# Mitochondrial Calcium Uptake Declines during Aging and is Directly Activated by Oleuropein to Boost Energy Metabolism and Skeletal Muscle Performance

**DOI:** 10.1101/2023.02.24.529830

**Authors:** Gaia Gherardi, Anna Weiser, Flavien Bermont, Eugenia Migliavacca, Benjamin Brinon, Guillaume E. Jacot, Aurélie Hermant, Mattia Sturlese, Leonardo Nogara, Denis Barron, Stefano Moro, Bert Blaauw, Rosario Rizzuto, Jerome N. Feige, Cristina Mammucari, Umberto De Marchi

## Abstract

Mitochondrial calcium (mtCa^2+^) uptake via the Mitochondrial Calcium Uniporter (MCU) couples the regulation of calcium homeostasis to energy production. mtCa^2+^ uptake is rate-limiting for mitochondrial activation during muscle contraction, but how MCU is affected during physiopathology and whether it can be stimulated therapeutically remains largely uncharacterized. By profiling human and preclinical aging of skeletal muscle, we discovered a conserved down-regulation of MCUR1 during aging that decreases mtCa^2+^ uptake and drives sarcopenia. Through a screen of 5000 bioactive nutrients, we identify the natural polyphenol Oleuropein as a specific MCU activator that stimulates mitochondrial respiration via binding to MICU1. Oleuropein activates mtCa^2+^ uptake and oxidative energy metabolism to enhance endurance and limit fatigue in vivo both in young and aged. These effects of Oleuropein are mediated by an MCU-dependent mechanism in skeletal muscle as they are lost upon muscle-specific MCU KO. Our work demonstrates that impaired mtCa^2+^ uptake causes mitochondrial dysfunction during aging and establishes Oleuropein as a novel nutrient that specifically targets MCU to stimulate mitochondrial bioenergetics and muscle performance.

## INTRODUCTION

Cellular energy production relies in large part on oxidative metabolism and ATP synthesis in mitochondria. Mitochondrial dysfunction is a major hallmark of aging that contributes to chronic diseases (Lopez-Otin et al., 2023; Sun et al., 2016), and an area of active therapeutic investigation. The anti-diabetic drug Metformin is being clinically tested to target aging given its geroprotective effects on lifespan and healthspan that acts at least in part by triggering mitochondrial adaptations via mitohormesis (Kulkarni et al., 2020). Other clinical therapeutic molecules such as NAD+ precursors (Covarrubias et al., 2021; Lapatto et al., 2023; Martens et al., 2018; Yoshino et al., 2021) autophagy/mitophagy activators (Andreux et al., 2019; Eisenberg et al., 2016; Ryu et al., 2016) or mitochondrial complex stabilizers (Chavez et al., 2020; Karaa et al., 2018) can prevent age-related decline indirectly by targeting other hallmarks of aging that maintain mitochondrial integrity and rewire bioenergetics during aging. In contrast, there is no approved therapy to directly stimulate the core ATP-generating bioenergetic capacity of mitochondria.

Mitochondrial metabolism is particularly important in skeletal muscle where contraction during physical activity is an extremely demanding process that consumes massive amounts of cellular energy (Hargreaves and Spriet, 2020). Exercise maintains mitochondrial fitness in muscle during aging (Menshikova et al., 2006) and mitochondrial dysfunction is a major contributor to the age-related decline of skeletal muscle (Hood et al., 2019; Migliavacca et al., 2019; Petersen et al., 2003; Short et al., 2005). Low mitochondrial activity of all respiratory complexes directly contributes to low muscle mass, low muscle strength and low walking speed (Andreux et al., 2018; Migliavacca et al., 2019; Zane et al., 2017) the clinical features of sarcopenia that impact physical autonomy, quality of life and mortality (Cruz-Jentoft et al., 2020). The medical management of sarcopenia relies on lifestyle interventions with exercise and nutrition (Robinson et al., 2018). While recent large clinical studies have demonstrated the benefits of structured exercise programs (Bernabei et al., 2022; Cartee et al., 2016; Pahor et al., 2014), nutritional management of sarcopenia largely relies on restoring the nutritional deficits of older adults in protein and micronutrients (Cruz-Jentoft et al., 2020; Deutz et al., 2014; Morley et al., 2010). Polyphenol-rich diets including Mediterranean diets rich in olive oil have also been associated with lower risk of chronic diseases of aging, in particular cardiovascular diseases (Trichopoulou et al., 2003; Trichopoulou et al., 1995). However, the nature and mechanisms of the individual polyphenols mediating these health benefits remain elusive, and the role of polyphenols in sarcopenia remains to be explored.

The regulation of mitochondrial Ca^2+^ (mtCa^2+^) directly controls oxidative metabolism and regulates skeletal muscle physiology (Gherardi et al., 2019; Rizzuto et al., 2012). The uptake of calcium in mitochondria is controlled by the Mitochondrial Ca^2+^ Uniporter (MCU), a multi-proteic complex composed of the MCU channel that transports Ca^2+^ across the inner mitochondrial membrane and of regulatory subunits that regulate MCU activity to adapt to the metabolic requirements of the cell (Baughman et al., 2011; De Stefani et al., 2011; Gherardi et al., 2020). During excitation-contraction coupling, Ca^2+^ is released from the sarcoplasmic reticulum to stimulate contraction and mtCa^2+^ couples the metabolic needs of contraction to ATP production via mitochondrial oxidative metabolism. Mechanistically, this is controlled by the activation of dehydrogenases of the TCA cycle that are acutely regulated via mtCa^2+^ levels. In particular, activation of the rate-limiting pyruvate dehydrogenase (PDH) occurs upon dephosphorylation by the Ca^2+^-dependent PDH phosphatase 1 (PDP).

Increased mtCa^2+^ and MCU activity in skeletal muscle have recently been linked to mitochondrial adaptations in response to different types of exercise training in humans (Zanou et al., 2021). Stimulating mtCa^2+^ uptake is sufficient to drive some of these adaptations as skeletal muscle over-expression of MCU in mice also promotes mitochondrial metabolism and structural muscle adaptations (Mammucari et al., 2015). Conversely, mtCa^2+^ uptake is rate limiting for normal muscle metabolism and contractile function as genetic deletion of MCU in skeletal muscle impairs mitochondrial oxidative capacity and reduces muscle force and exercise performance (Gherardi et al., 2019; Mammucari et al., 2015). In addition, regulatory subunits of the MCU complex fine tune muscle physiology as skeletal muscle-specific deletion of MICU1 compromises mtCa^2+^ uptake and energy metabolism, leading to myofiber damage and muscle weakness and fatigue (Debattisti et al., 2019). Human genetics have also confirmed a central role of mtCa^2+^ uptake via the MCU complex in muscle physiology as monogenic mutations of MICU1 cause rare orphan diseases with myopathic phenotypes (Logan et al., 2014), that present with severe fatigue and lethargy (Lewis-Smith et al., 2016). Although these studies collectively highlight a robust link between mtCa^2+^ regulation and skeletal muscle health, the role of MCU in chronic conditions such as muscle wasting disorders and aging is not well understood, and translatable therapeutic tools and interventions to boost mtCa^2+^ by activating MCU are lacking.

Here we discovered that the capacity for mtCa^2+^ uptake declines during aging in model organisms and in humans via downregulation of MCUR1, and directly impairs mitochondrial energy metabolism. To overcome this, we screened 5000 natural bioactive molecules present in food and discovered the oleuropein olive polyphenol as a potent and selective activator of MCU with therapeutic potential for skeletal muscle. Oleuropein stimulates mtCa^2+^ uptake via direct binding of its deglycosylated metabolite to the MICU1 subunit of MCU, inducing acute ergogenic effects and chronic adaptations to increase oxidative metabolism and muscle performance *in vivo*. Oleuropein also overcomes the age-related MCUR1-dependent impairment of mtCa^2+^ and rescues mitochondrial activity and physical performance in aged mice. Our work establishes the regulation of MCU-dependent mtCa^2+^ uptake as a novel mechanism of aging, and demonstrates that oleuropein is a first-in-class natural molecule to enhance muscle performance and prevent muscle aging by directly activating MCU and mitochondrial bioenergetics with high specificity.

## RESULTS

### Mitochondrial Ca^2+^ uptake declines during aging in human skeletal muscle via downregulation of MCUR1

Given the regulation of mitochondrial bioenergetics by mtCa^2+^ and the role of mitochondrial dysfunction in driving aging of skeletal muscle, we investigated whether mtCa^2+^ participates to the physiopathology of aging muscle. The mtCa^2+^ profile of primary human myotubes (**Fig. 1A**) from adult and old donors was compared by stimulating the ryanodine receptor (RyR1) with caffeine to mimic contraction-induced Ca^2+^ release from the sarcoplasmic reticulum (SR). The capacity of skeletal muscle mitochondria to take up Ca^2+^ was strongly impaired by aging as mtCa^2+^ uptake was reduced by 45% in aged individuals (**Figs. 1B-1C**).

**FIGURE 1.**
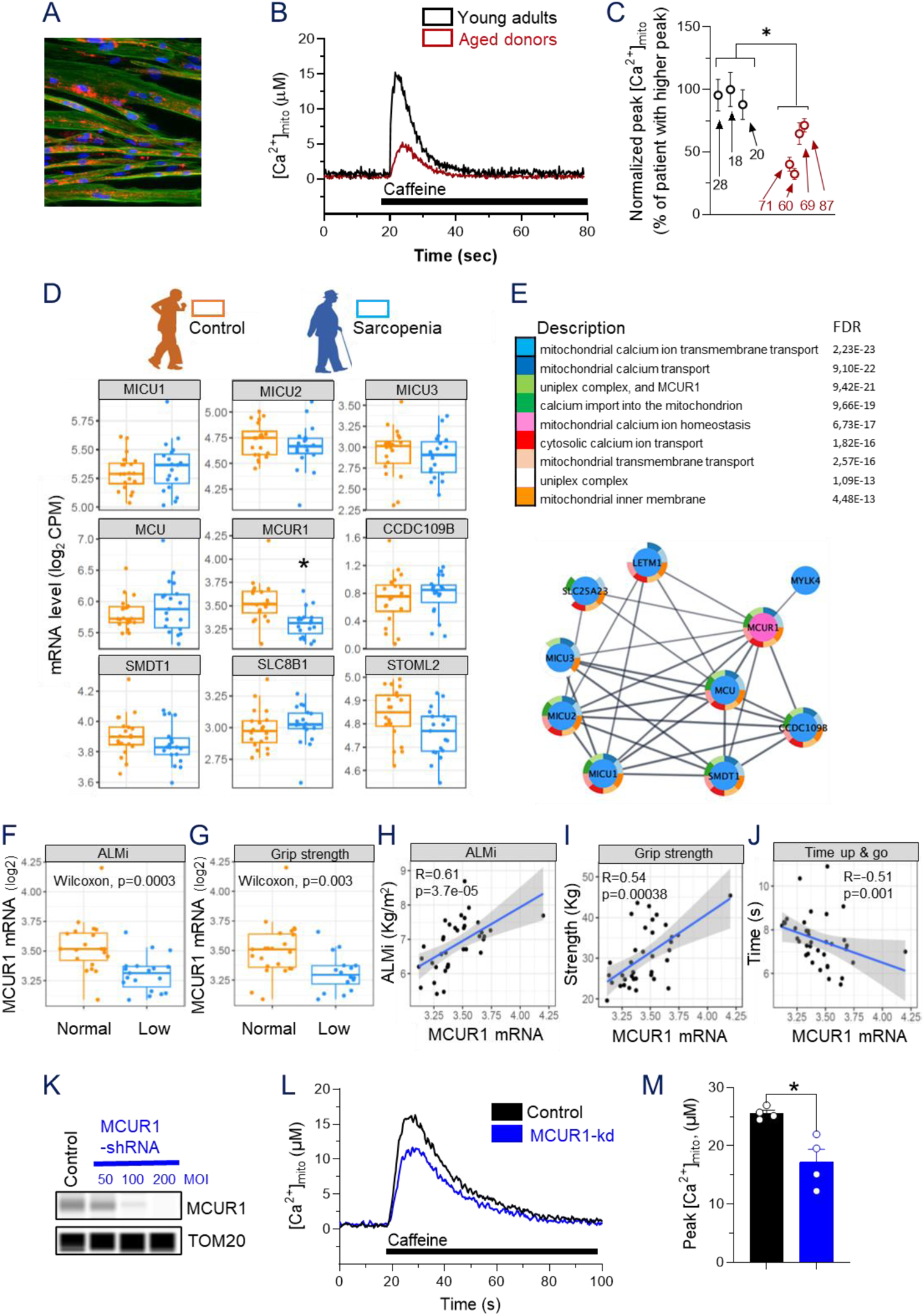
Uptake of mitochondrial Ca2+ declines during aging in human skeletal muscle. (A-C) Aequorin-based mtCa^2+^ uptake in primary human myotubes from aged vs young donors. (A) Primary human myoblasts differentiated in myotubes and immunostained for fibers (troponin-t, green), mitochondria (MCU, red) and nuclei (DAPI, blue). Representative mtCa^2+^ traces (B) and quantification of the mtCa^2+^ uptake peak (C) in myotubes from young and aged donors (n>5 samples/donor). (D) mRNA expression of genes regulating mtCa^2+^ measured by RNA sequencing of human skeletal muscle biopsies of sarcopenic vs. age-matched healthy older people (n=19-20 muscle samples/group; CPM: counts per million). (E) Gene-set enrichment analysis of skeletal muscle genes associated with MCUR1 expression. (Upper panel) Gene sets are ordered according to the significance of their association with MCUR1. (Lower panel) Interactome map of MCUR1-linked proteins. (F, G) MCUR1 mRNA expression in skeletal muscle biopsies of older people with low appendicular lean mass index (*ALMI,* F) or low grip strength (G). (H-J) Correlation of MCUR1 mRNA expression in skeletal muscle biopsies of older people with with ALMI (H), grip strength (I), and performance in the “time up and go” test (J). (K-M) MCUR1 knockdown (MCUR1-kd) decreases mtCa^2+^ uptake during caffeine stimulation. (K) Validation of MCUR1 protein expression by capillary western blot after MCUR1-kd using adenoviral infection-mediated shRNA delivery in primary human myotubes. Scramble shRNA was used as control and TOM20 was used as internal standard of mitochondrial content. MOI: multiplicity of infection. (L) Representative mtCa^2+^ traces after MCUR1-kd in primary human myotubes. (M) Quantification of the effect of MCUR1-kd on mtCa^2+^ uptake peak, after caffeine stimulation. In all bar graphs, data are presented as mean ± SEM with *p < 0.05 using a two-tailed unpaired T-test.

To characterize the molecular mechanisms through which skeletal muscle aging impairs mtCa^2+^, we analyzed the expression of genes controlling mtCa^2+^ uptake and extrusion in human muscle biopsies from older people with sarcopenia (Migliavacca et al., 2019). Sarcopenia significantly reduced the expression of MCUR1 (**Fig. 1D**), a regulator of mtCa^2+^ transport (Chaudhuri et al., 2016; Mallilankaraman et al., 2012; Paupe et al., 2015; Tomar et al., 2016; Vais et al., 2015), while other genes regulating mtCa^2+^ were not altered. Pathway and interactome analyses confirmed that decreased MCUR1 expression in sarcopenia controls a molecular node regulating mtCa^2+^ homeostasis via the MCU (**Fig. 1E**). Downregulation of MCUR1 was linked to both low muscle mass assessed via appendicular lean mass index (ALMi) by DXA (**Fig. 1F**) and muscle strength assessed by grip strength (**Fig. 1G**). In continuous variable analysis, the muscle expression of MCUR1 also positively associated with ALMi, grip strength and gait speed (**Fig. 1H-I & S1A**), and negatively associated with the time required to perform a “Time up and go” or chaise rise test (**Figs. 1J & S1B**), all indicative that MCUR1 expression is high in elderly people with high physical capacity. These results demonstrate that the regulation of mtCa^2+^ transport is altered in human skeletal muscle during aging and physical decline, and suggest a possible role of MCUR1 in mediating the age-related impairment of mtCa^2+^ uptake.

To test if the down-regulation of MCUR1 observed during sarcopenia is sufficient to impair skeletal muscle mtCa^2+^ uptake, we depleted MCUR1 in primary human myotubes from young adults using viral shRNA over-expression (**Fig. 1K**). MCUR1 knock-down reduced mtCa^2+^ uptake after caffeine stimulation by 33% (**Figs. 1L-M**), demonstrating that the downregulation of MCUR1 is sufficient to recapitulate the effects of aging on mtCa^2+^ uptake (**Figs. 1B-1C**). Taken together, these results demonstrate a functional link between impaired mtCa^2+^ activation and muscle decline during aging that is mediated by the down-regulation of MCUR1.

### Impaired mitochondrial Ca^2+^ uptake alters pyruvate dehydrogenase (PDH) activity and energy metabolism in a pre-clinical model of natural aging

To investigate whether the regulation of mtCa^2+^ homeostasis occurring during human aging can be modelled preclinically, we analyzed mtCa^2+^ uptake and expression of genes regulating mtCa^2+^ uptake in the skeletal muscle of mice aged 24 months compared to young controls. As previously reported (Weisleder et al., 2006), cytosolic Ca^2+^ transients generated upon caffeine stimulation were decreased in aged myofibres while resting cytosolic Ca^2+^ levels were unaltered **(Fig. S2A, S2B)**. MtCa^2+^ uptake measured in isolated mouse myofibers upon caffeine-induced Ca^2+^ release decreased during aging **(Figs. 2A-2B)**, while resting mtCa^2+^ levels were unaltered **(Fig. 2C)**. Like in human muscle aging, the regulation of mtCa^2+^ uptake directly contributed to this phenotype as MCUR1 was downregulated by 54% during natural mouse aging, while the expression of the channel (MCU, MCUb and EMRE) and of other regulatory subunits (MICU1, MICU1.1, and MICU2) were unaltered in aged mice **(Figs. 2D & S2C)**. As schematized in **Fig. 2E**, mitochondrial matrix Ca^2+^ acutely activates pyruvate dehydrogenase (PDH) via dephosphorylation mediated by allosteric activation of the Ca^2+^-dependent PDH phosphatase (PDP). We previously demonstrated that decreased mtCa^2+^ uptake triggered by a muscle-specific deletion of MCU (skMCU^−/−^) reduces PDH activity, impairs mitochondrial oxygen consumption rate (OCR), and rewires substrate dependency from glucose to fatty acid oxidation (Gherardi et al., 2019), all in a PDP-dependent fashion as PDP overexpression is sufficient to restore these phenotypes (Gherardi et al., 2019). We hypothesized that the reduction in mtCa^2+^ uptake observed during skeletal muscle aging could trigger similar metabolic impairments. PDH phosphorylation levels, which inversely correlate with PDH activity, increased by 54% in aged muscle **(Fig. 2F-G)**, and basal, ATP-dependent and maximal OCR measured in isolated myofibers declined in the muscle of aged mice **(Fig. 2H-I)**. Inhibition of fatty acid oxidation with etomoxir had a greater impact on OCR in aged muscle, demonstrating that aging shifts the substrate preference of mitochondria towards FA oxidation **(Figs. 2J-K)**.

**FIGURE 2.**
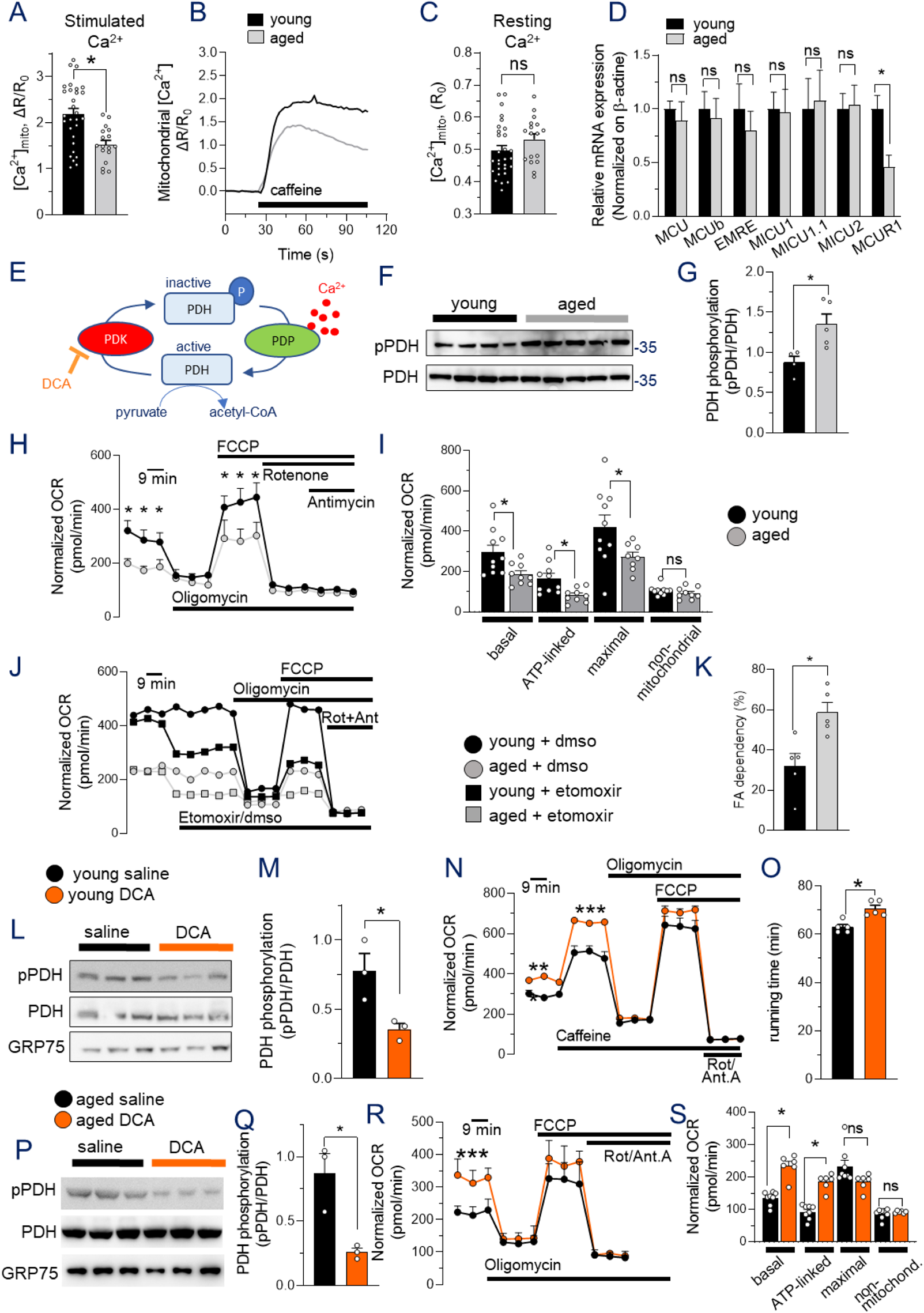
Impaired mitochondrial Ca^2+^ uptake during aging impairs energy metabolism via PDH in skeletal muscle. (A-C) Mt-fura-2-based mtCa^2+^ measurement of flexor digitorum brevis (FDB) myofibers from aged vs young mice. Quantification (A) and representative traces (B) of peak mtCa^2+^ uptake upon caffeine-stimulation, and resting mtCa^2+^ levels (C) (n>20 fibers from 3 mice / group). (D) Relative mRNA expression levels of MCU complex genes in tibialis anterior (TA) muscles from aged vs young mice (n=5 mice/group). (E) Representative scheme of PDH regulation. Ca^2+^ binding in the mitochondrial matrix activates pyruvate dehydrogenase phosphatase 1 (PDP1), the Ca^2+-^dependent isoform of the PDP enzyme that, in turn, dephosphorylates and activates PDH, leading to acetyl-CoA production. Dichloroacetate (DCA) inhibits PDK. (F-G) PDH phosphorylation levels quantified by western blot in aged TA muscles from aged vs young mice (n=4-5 mice/group). (H-I) Oxygen consumption rate (OCR) in isolated FDB myofibers from aged vs young mice. Data are normalized on mean calcein fluorescence (H) and components of respiration accounting for basal, ATP-dependent (quantified after oligomycin treatment), maximal (quantified after FCCP treatment) and non-mitochondrial (quantified after rotenone + antimycin treatment) respiration are quantified in (I). (n=10 samples/group). (J-K) Fatty acid (FA) dependent-respiration measured in FDB myofibers from aged vs young mice. Etomoxir was used to inhibit FA utilization. FA dependency was calculated and expressed as the etomoxir-sensitive percentage of basal OCR (K). Data are normalized on mean calcein fluorescence (n=5 samples/group). (L-S) Effects of DCA-mediated PDH activation in skeletal muscle of young (L-O) and aged (P-S) mice. (L-M & P-Q) PDH phosphorylation levels quantified by western blot in TA muscles of DCA-treated mice compared to saline controls using GRP75 as loading control; n=3 mice/group. (N;R-S) OCR measured in FDB myofibers and normalized on mean calcein fluorescence in DCA-treated mice compared to saline controls (n=6-10 samples/group). (O) Exercise performance assessed by maximal treadmill running time in DCA-treated mice compared to saline controls. (n=5 mice/group). In all bar graphs, data are presented as mean ± SEM with *p < 0.05 using a two-tailed unpaired T-test.

Pharmacological activation of the PDH complex by dichloroacetate (DCA) **(Fig. 2E)** has previously been used to increase oxidative phosphorylation activity and muscle strength in a mouse model of amyotrophic lateral sclerosis (Palamiuc et al., 2015). We used in vivo DCA treatment to test if boosting PDH activity could also stimulate metabolism and performance of young healthy mice or rescue the consequences of impaired genetic or age-related mtCa^2+^ uptake. In young mice, DCA treatment stimulated muscle PDH by reducing its phosphorylation **(Figs. 2L-M)**, increased myofiber respiration **(Fig. 2N)**, and boosted endurance exercise performance **(Fig. 2O)**. DCA could overcome impaired mitochondrial bioenergetics caused by genetically altered mtCa^2+^ uptake as OCR increased and FA dependency decreased in skMCU−/− myofibres treated with DCA **(Fig. S2D-E)**. Finally, DCA treatment in aged animals decreased PDH phosphorylation **(Figs. 2P-Q)** and increased basal OCR **(Figs. 2R-S)**. These results indicate that the MCU-PDH axis declines in aged muscle and that restoring PDH activity overcomes the bioenergetic defects caused by impaired mtCa^2+^ uptake during aging.

### A high throughput screen identifies the natural polyphenol Oleuropein as a potent activator of mitochondrial calcium uptake

Given the link of mtCa^2+^ uptake with mitochondrial energy metabolism and muscle performance and its impairment during skeletal muscle aging, we performed a high throughput (HT) screen of natural molecules that activate mtCa^2+^ to identify novel nutritional solutions that could stimulate mitochondrial bioenergetics and reverse age-related decline (**Figs. S3A-B**). We analyzed 5571 natural molecules present in food on mtCa^2+^ uptake in Hela cells stimulated with histamine (**Fig. 3A-B & S3C)**, using kaempferol as a positive control (Montero et al., 2004). 78 primary hits, corresponding to a rate of 1.4% positive hits (**Fig. 3A-B**), were filtered by orthogonal screening (**Fig. 3C**) and the resulting 52 positive hits were counter-screened for their ability to specifically increase mtCa^2+^ without triggering cytosolic Ca^2+^ rise (**Fig. 3D**). By filtering the 43 molecules that specifically and reproducibly activate mtCa^2+^ for known safety and bioavailability in humans, we identified that the olive polyphenol Oleuropein has the best profile of bioactive and safe mtCa^2+^ activation (**Fig. 3E).** We confirmed the effect of oleuropein in intact cells as a specific mitochondrial (**Figs 3F-G**), and not cytosolic Ca^2+^ activator (**Figs 3H-I**). In addition, oleuropein also increased mtCa^2+^ uptake in semi-permeabilized Hela cells stimulated with exogenous Ca^2+^ to over-ride Ca^2+^ release from the reticulum (**Figs 3J-K**), demonstrating that the effect of oleuropein on mtCa^2+^ transport is direct and independent of Ca^2+^ mobilization from endoplasmic reticulum. Since oleuropein is a glycosylated molecule that is metabolized upon intestinal absorption (**Fig. 3L**; (Clewell et al., 2016)), we also tested the ability of secondary oleuropein metabolites to promote mitochondrial calcium uptake in dose response (**Fig. 3M-O)**. Oleuropein aglycone (oleuropein-a) was the most potent metabolite and stimulated mtCa^2+^ uptake with an EC50 of 5μM (**Fig. 3N)**, while the secondary metabolite hydroxytyrosol could stimulate mtCa^2+^ uptake with weaker potency and affinity (**Fig. 3O)**. These results demonstrate that oleuropein and its direct metabolites are potent and direct activators of mtCa^2+^ uptake with a favorable safety and bioavailability for human clinical use.

**FIGURE 3.**
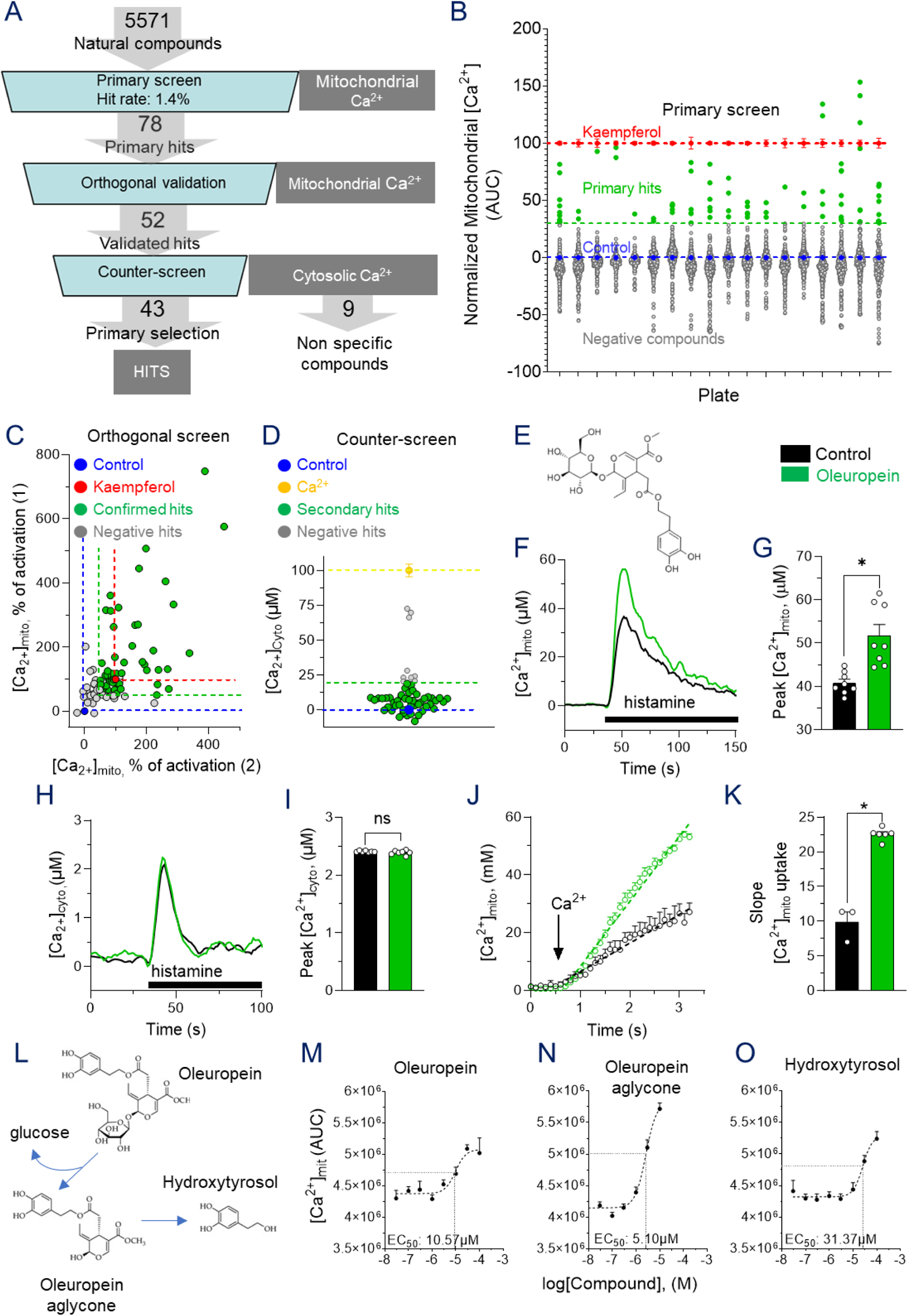
A high-throughput screen identifies the natural polyphenol Oleuropein as a potent activator of mitochondrial calcium uptake. (A-D) Aequorin-based, high-throughput assay of mtCa^2+^ uptake was developed to select MCU-specific natural bioactives from a library of 5571 natural food bioactive molecules. (A) Workflow and results of the screen. (B) Results of mtCa^2+^ primary screen in HeLa cells stimulated with 100 μM histamine; primary hits (green), non-active molecules (grey). Results are scaled normalized with negative control at 0 (blue) and positive control with 20μM kaempferol at 100 (red). (C) Orthogonal replication of positive hits from the primary screen using the same assay. Positive control, kaempferol 10μM. (D) Counter screen of cytosolic Ca^2+^ to exclude the hits that modulate both mtCa^2+^ and cytosolic Ca^2+^; positive control is 100mM extracellular Ca^2+^. For A-D, n=1 replicate/molecule. (E-I) Selection and validation of the positive hit oleuropein. (E) Chemical structure of the polyphenol oleuropein. (F-G) Effect of 10μM Oleuropein on mtCa^2+^ uptake in Hela cells stimulated with 100μM histamine, with quantification of mtCa^2+^ peak uptake from n=8 independent experiments (G). (H-I) Effect of 10μM Oleuropein on cytoplasmic Ca^2+^ in Hela cells stimulated with 100μM histamine, with quantification of cytosolic Ca^2+^ peak from n=7 independent experiments (I). (J-K) Effect of 10 μM Oleuropein aglycone on MCU activity in semi-permeabilized Hela cells evoked with 4 μM free Ca^2+^, with quantification of the slope of mtCa^2+^ uptake from n=3-6 cell culture replicates/group (K). (L-O) Effect of Oleuropein and its metabolites, oleuropein aglycone and hydroxytyrosol, on - mtCa^2+^ uptake induced by caffeine in C2C12-derived myotubes. Structural formula of the molecules (L) and dose-response effect of oleuropein (M), oleuropein aglycone (N) and hydroxytyrosol (O) from n=6-8 cell culture replicates per molecule. In all bar graphs, data are presented as mean ± SEM with *p < 0.05 using a two-tailed unpaired T-test.

### Oleuropein requires MICU1 to stimulate MCU-dependent mitochondrial energy metabolism in primary human myotubes

Given the role of mtCa^2+^ in coupling contraction with energy production in skeletal muscle, we investigated if oleuropein also enhances mtCa^2+^ and energy metabolism in models of human skeletal muscle and dissected how its primary circulating metabolite oleuropein aglycone (oleuropein-a) can stimulate mitochondrial calcium uptake at the molecular level in muscle cells. Oleuropein-a potently stimulated mtCa^2+^ uptake in primary myotubes differentiated from human skeletal muscle and stimulated with caffeine to mimic contraction (**Figs. 4A**), without affecting cytosolic Ca^2+^ elevation (**Figs. S4A-B)**. Similar to what was observed in HeLa cells, the primary circulating oleuropein aglycone metabolite was more active than the parent glycosylated molecule and the downstream metabolite hydroxytyrosol to raise mtCa^2+^ in human muscle cells (**Fig. 4B**), prompting to focus further cellular exploration on deglycosylated oleuropein. Consistent with the rate limiting role of MCU for mtCa^2+^ uptake (Baughman et al., 2011; De Stefani et al., 2011), AAV-mediated shRNA knockdown of MCU (MCU-kd) that strongly reduced MCU protein levels abolished caffeine-induced mtCa^2+^ uptake (**Fig. 4C**). Importantly, MCU-kd completely inhibited the ability of oleuropein to stimulate mtCa^2+^ uptake (**Figs. 4D-E**), demonstrating that oleuropein requires the channel subunit of the MCU complex to stimulate mtCa^2+^ uptake.

**FIGURE 4.**
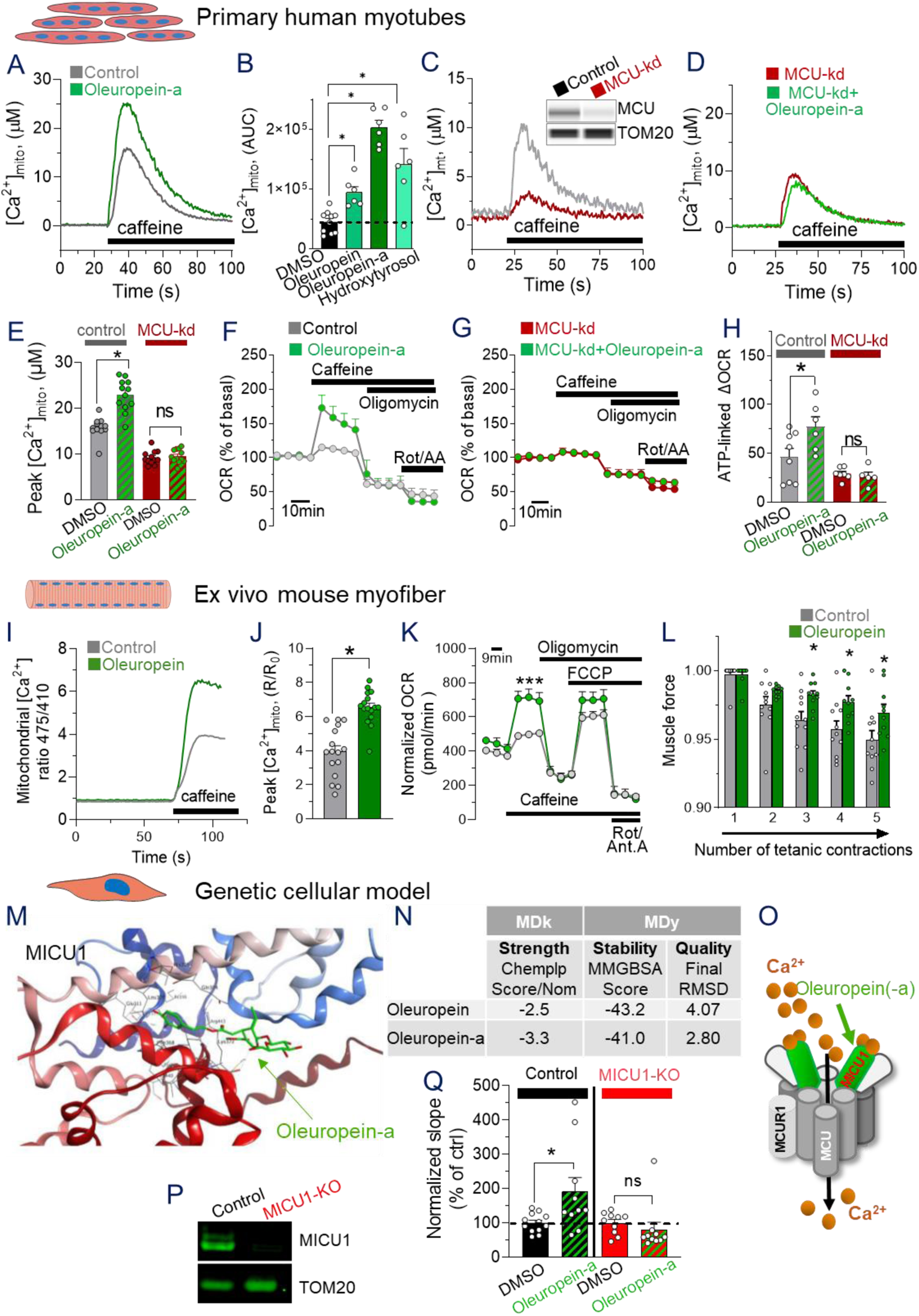
Oleuropein activates mitochondrial energy metabolism in primary human myotubes and intact skeletal muscle via MICU1-dependent mtCa^2+^ uptake. (A-E) Aequorin-based mtCa^2+^ uptake in control (A-B;E) and MCU knock-down (C-E) primary human myotubes treated with 10μM Oleuropein metabolites. (C) Caffeine stimulated-mtCa^2+^ peak was quantified from n=6-12 cell culture replicates / group. (F-H) Oxygen consumption rate (OCR) of control (F;H) and MCU knock-down (G-H) primary human myotubes treated with 10μM Oleuropein-a. The Ca^2+^-dependent mitochondrial respiratory capacity for ATP production was calculated as the OCR difference (ΔOCR) between caffeine- and oligomycin-dependent respiration in n=6-12 cell culture replicates / group (H). (I-J) 4mtGCaMP6f-based mtCa^2+^ uptake in isolated mouse FDB fibers treated with 10μM Oleuropein. Caffeine stimulated-mtCa^2+^ peak was quantified from n>20 fibers/group. (K) OCR of isolated FDB myofibers treated with 10μM Oleuropein. Data are normalized on mean calcein fluorescence (n=10 samples/ group). (L) Ex vivo fatigue of mouse EDL muscle treated with 10μM oleuropein during repeated electrical stimulation of tetanic force; n=10 mice/group. (M-Q) Oleuropein activates mtCa^2+^ uptake by binding MICU1. (M) In silico docking of Oleuropein-a (green) to the MICU1 crystal structure. (N) Quantification of Molecular Docking (MDk) and Molecular Dynamics (MDy) of oleuropein and oleuropein-a to MICU1. (O) Model of oleuropein-dependent activation of mtCa^2+^ uptake via MICU1 binding. (P) Western blot validation of MICU1-KO in MEF cells with TOM20 as loading control. (Q) MCU activity in semi-permeabilized control and MICU1-KO MEF cells treated with 10 μM Oleuropein-a, with quantification of the slope of mtCa^2+^ uptake evoked with 4 μM free Ca^2+^ from n=10 cellular replicates. In all bar graphs, data are presented as mean ± SEM. *p < 0.05 and ns: not significant using a one-way ANOVA (A-H) or a two-tailed unpaired T-test (J-L;Q).

Using oxygen consumption in primary human myotubes stimulated by caffeine as a model of contraction-induced mitochondrial energy metabolism, we demonstrated that Oleuropein-a acutely increases mitochondrial respiration by +67% during caffeine-induced Ca^2+^ mobilization and boosts the functional generation of ATP via an oligomycin-sensitive coupled respiration (**Fig. 4F-H**). In line with the results on mtCa^2+^ (**Fig. 4E**), knockdown of MCU completely prevented the effects of oleuropein-a on coupled respiration (**Figs. 4G and 4H**), demonstrating that MCU and mtCa^2+^ are required for oleuropein-dependent mitochondrial metabolism. We next measured the effect of oleuropein-a on cell death, to verify that the augmented mtCa^2+^ uptake induced by oleuropein remains physiological. Consistent with published data from literature on the safe properties of Oleuropein (Clewell et al., 2016; Guex et al., 2018), and on the effects of MCU overexpression in skeletal muscle (Mammucari et al., 2015), oleuropein and its metabolites did not affect cell death in primary human myotubes (**Fig. S4C**).

In order to demonstrate that Oleuropein can activate mtCa^2+^ uptake in fully functional muscle fibers and investigate the metabolic consequences of this activation on contractile properties, we treated isolated myofibers from mouse flexor digitorum brevis (FDB) muscle *ex vivo*. In this system, Oleuropein and its metabolites also enhanced caffeine-induced mtCa^2+^ uptake (**Figs. 4I-J; S4D),** and mitochondrial respiratory capacity (**Fig. 4K**). To determine the effect of Oleuropein on skeletal muscle performance, we measured muscle strength and fatigue. Oleuropein did not show any detrimental effect on force production (**Fig. S5**). Mitochondrial metabolism and oxidative capacity, that are boosted by oleuropein (**Figs. 4F,4H,4K**), are important to sustain muscle energy production during repeated contractions and to prevent fatigue during physical activity. Oleuropein-treated myofibers could maintain high intensity force during repeated contractions better than the untreated controls, demonstrating that oleuropein lowers fatiguability during exercise (**Fig. 4L)**. Altogether, these results demonstrate that oleuropein enhances mitochondrial energy metabolism across different models of skeletal muscle and improves the resistance to fatigue during contraction. We then performed in silico docking simulations to the crystal structure of MCU subunits in order to understand how oleuropein activates the MCU complex. Given the optimal position of MICU1 for interactions with mitochondrial intermembrane molecules that diffuse from the cytoplasm, and its ability to interact with ligands (Di Marco et al., 2020), we tested if oleuropein could activate MCU via interaction with MICU1. Through molecular docking and dynamics, we identified that Oleuropein and Oleuropein-a can bind a cleft of human MICU1 (**Fig. 4M-N),** that was also recently reported as a ligand binding pocket (Di Marco et al., 2020). For the analysis of the docking, we measured the strength of the interaction (**Fig. 4N**). The stability of conformation and the quality of the interaction indicated that both the compounds are well accommodated into the binding site. Here we report the best conformation obtained for oleuropein aglycone (**Fig. 4M**). Oleuropein and Oleuropein-a were similarly oriented in the MICU1 binding site with the hydroxytyrosol portion of the molecule binding Glu311 of MICU1 and the elenolic group binding both the MICU1 N-lobe (216-220) and C-lobe (loop 441-443) (**Fig. S6** and **Movie S1**).

The functional relevance of the in silico binding was confirmed experimentally through a loss of function of MICU1 using MICU1 KO MEF cells (Antony et al., 2016), where mtCa^2+^ uptake was measured in Ca^2+^-stimulated semi-permeabilized cells. Similar to other models, oleuropein efficiently stimulated mtCa^2+^ uptake in WT cells, but this response was completely blunted in MICU1 KO cells (**Fig. 4P-Q**). This cellular and in silico experiments demonstrate that oleuropein activates mtCa^2+^ uptake via an MCU-dependent mechanism that relies on direct binding to MICU1 (**Fig. 4O**).

### Acute and chronic dietary treatment with Oleuropein enhances muscle energy metabolism and performance via MCU

To determine the impact of Oleuropein on energy metabolism and muscle performance in a complex physiological organism, we treated young adult mice orally via dietary administration of an olive leaf extract (OLE) standardized for high oleuropein content. A single oral dose of OLE acutely increased mtCa^2+^ uptake measured *ex vivo* in isolated myofibers (**Figs. 5A and 5B**). In a 2 to 8 hour time course after acute oral intake, OLE induced a sustained 36-58% dephosphorylation of PDH in skeletal muscle that was already prominent after 2h, consistent with the reported rapid peak of oleuropein and its metabolites in blood (Garcia-Villalba et al., 2014; Polia et al., 2022) (**Fig 5C and S7A**). As expected, dephosphorylation was mirrored by increased PDH activity in skeletal muscle which was boosted by +44% after acute OLE intake (**Fig. 5D**). Consistent with PDH activation which is rate limiting for pyruvate oxidation and mitochondrial oxidative metabolism, *in vivo* OLE treatment also stimulated oxygen consumption in myofibers (**Fig. S7B**). The effects of OLE in muscle were fully maintained after chronic treatment as dietary administration of OLE for 1 month in adult mice increased mtCa^2+^ uptake (**Figs. 5E and 5F**), decreased PDH phosphorylation (**Fig. 5G and S7C**), increased PDH activity (**Fig. 5H**), and stimulated oxygen consumption and mitochondrial activity (**Fig. 5I**). Muscle mass was not affected by OLE in young adult mice (**Fig S7D**). In contrast, OLE enhanced muscle performance by improving the resistance to fatigue in an electrically stimulated muscle force assay (**Fig 5J**), confirming the results obtained in isolated myofibers (**Fig. 4L**). In addition, OLE enhanced exercise performance in a treadmill endurance test **(Fig. 5K**). Since polyphenols can have pleiotropic effects and oleuropein has been reported to act via distinct mechanisms including as an anti-oxidant (Bulotta et al., 2014), we tested whether the effects of OLE on mitochondrial bioenergetics and muscle performance are mediated by MCU. To this end, dietary OLE was administered chronically to mice with a skeletal muscle-specific deletion of MCU using Cre/lox recombination under control of the myosin light chain 1c promoter (skMCU^−/−^ mice) (Gherardi et al., 2019). Unlike in WT (**Figs. 5I and K)** and control (**Figs. 5L-M & S7E)**, OLE had no effect on PDH phosphorylation mitochondrial oxygen consumption and treadmill performance in skMCU^−/−^ mice (**Figs. 5N-O; Fig. S7F)**. These results demonstrate that the stimulation of mitochondrial calcium uptake by OLE in healthy adults acutely and chronically activates PDH, stimulates mitochondrial respiration and enhances exercise performance specifically via MCU.

**FIGURE 5.**
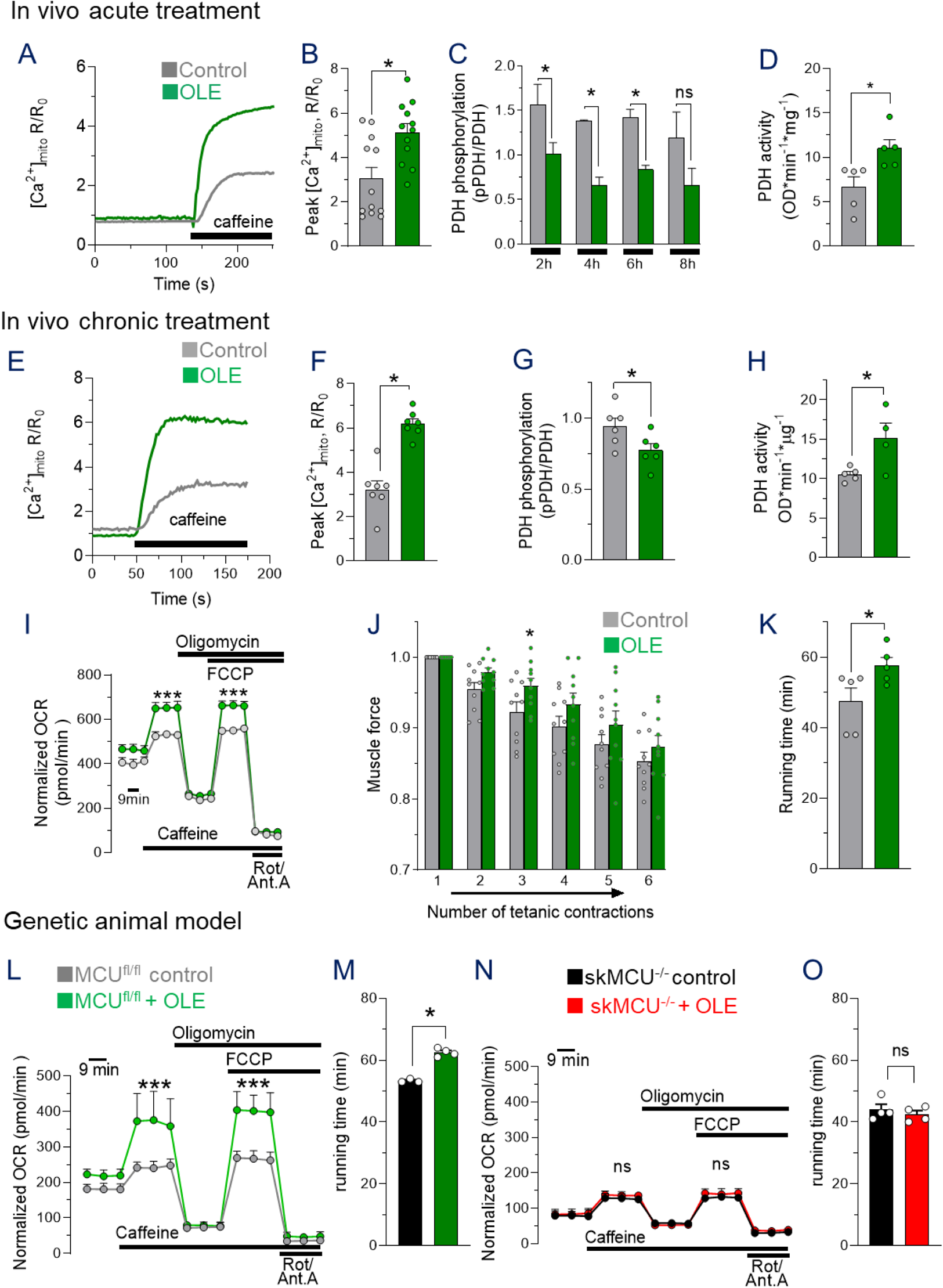
Acute and chronic dietary Oleuropein activates skeletal muscle energy metabolism and enhances performance in vivo via MCU. (A-D) 4mtGCaMP6f-based mtCa^2+^ uptake (A-B), PDH phosphorylation (C) and PDH activity (D) in skeletal muscle of young adult mice acutely receiving a single oral dose of olive leaf extract with 40% oleuropein (OLE) at 50mg/kg or control vehicle. Representative mtCa^2+^ traces (A) and quantification of mtCa^2+^ peaks (B) in caffeine-stimulated isolated FDB myofibers of mice acutely treated with OLE (n=12 fibers from different mice / group). PDH activity was measured in Tibialis Anterior muscle via PDH dephosphorylation by western blot 2-8 hours after oral OLE (C; n=2 mice/group) and PDH enzymatic activity 6h after oral OLE (D; n=5 mice/group). (E-K) 4mtGCaMP6f-based mtCa^2+^ uptake (E-F), PDH phosphorylation (G), PDH activity (H), mitochondrial respiration (I) and muscle performance (J-K) in young adult mice fed a control or OLE diet equivalent to 50mg/kg/day for 1 month. Representative mtCa^2+^ traces (E) and quantification of mtCa^2+^ peaks (F) in caffeine-stimulated isolated FDB myofibers of mice chronically receiving OLE diet (n=7 fibers/group). PDH activity in Tibialis Anterior muscle measured via PDH dephosphorylation normalized to GRP75 by western blot (G) and PDH enzymatic activity (H) of mice fed for 1 month with OLE diet (n=4-6 mice/group). (I) Mitochondrial respiratory capacity measured via oxygen consumption rate (OCR) normalized to mean calcein fluorescence in isolated FDB myofibers of mice fed for 1 month with OLE diet (n=10 samples/ group). (J) In vivo fatigue of gastrocnemius muscle during repeated electrical stimulation of tetanic force in mice fed for 1 month with OLE diet; n=10 mice/group. (K) Exercise performance assessed in mice fed for 2 weeks with OLE diet. (n=5mice/group). (L-O) Mitochondrial respiration after 4 weeks (L;N) and exercise performance after 2 weeks (M;O) of feeding with control or OLE diet equivalent to 50mg/kg/day in young control (MCU^fl/fl^; L-M) or skeletal-muscle specific MCU KO (skMCU^−/−^; N-O) mice (n=3-4mice/group). In all bar graphs, data are presented as mean ± SEM with *p < 0.05 using a two-tailed unpaired T-test.

### Oleuropein reverses the age-related decline of muscle bioenergetics and performance across species

Building on the observation that mtCa^2+^ uptake and mitochondrial metabolism decline during aging and sarcopenia via a MCUR1-driven mechanism (**Fig. 1A-J**; (Migliavacca et al., 2019)), we tested whether oleuropein binding via MICU1 could still activate mtCa^2+^ uptake in aged muscle with decreased MCUR1. The effect of oleuropein on mtCa^2+^ uptake was dominant over impaired muscle MCU activity caused by downregulation of MCUR1 (**Fig. 1K-M**) as oleuropein-a could efficiently stimulate mtCa^2+^ uptake in human primary myotubes where MCUR1 was genetically knocked-down (**Figs. 6A-B & 1K**). Oleuropein-a also reversed age-related decline of mtCa^2+^ uptake in primary myotubes from aged human donors (**Figs. 6C-D**). Since the decline of MCUR1 and mtCa^2+^ occurring during human aging is conserved in rodent models (**Fig. 2A-G**), we used complementary in vivo models to test if stimulating mtCa^2+^ uptake with oleuropein can reverse age related muscle decline. In a rat model of sarcopenia with strong mitochondrial dysfunction (Ibebunjo et al., 2013; Pannerec et al., 2016), chronic dietary OLE administration after the onset of sarcopenia robustly reduced PDH phosphorylation by 60% (**Fig. 6E-G)**, indicative of enhanced activity and restored mitochondrial oxidative metabolism. Chronic dietary OLE also enhanced mitochondrial respiration of aged myofibers (**Fig. 6H**) and restored exercise performance (**Fig. 6I**) in a mouse aging model where mitochondrial activity and running performance decline (**Fig. 2H** and (Yanai and Endo, 2021)). In addition, and distinctly from young adults (**Fig S7D**), OLE had a beneficial effect on muscle weight (**Fig. 6J**) in aged mice.

**FIGURE 6.**
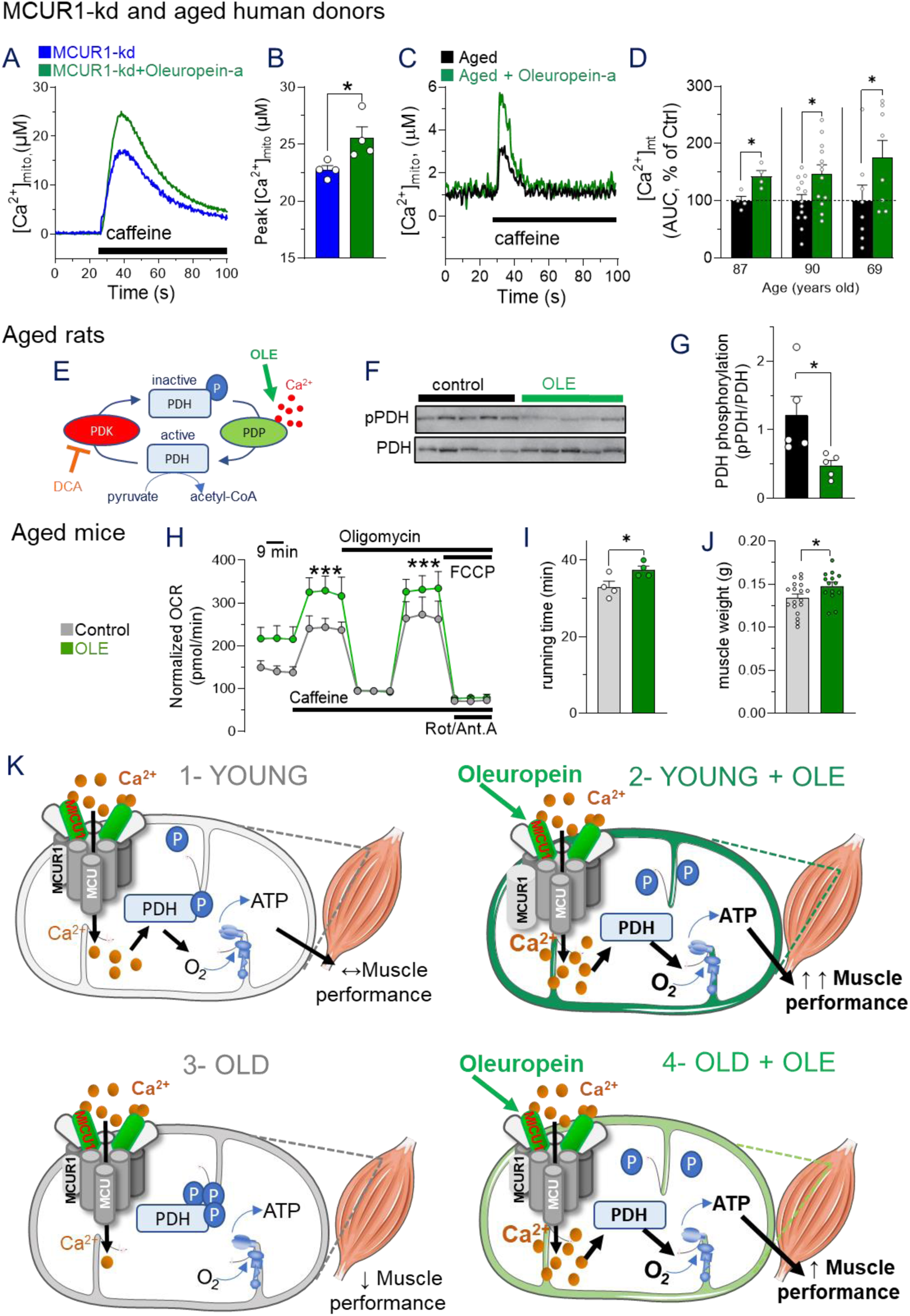
Oleuropein reverses age-related decline of mtCa^2+^ uptake, energy metabolism and muscle performance across species. (A-D) mtCa^2+^ uptake in primary human myotubes with MCUR1 KD (A-B) or from aged donors (C-D) and treated with 10μM oleuropein-a; n=4 or more cellular replicates/condition. (E-G) PDH activation measured by dephosphorylation by western blot relative to GRP75 in gastrocnemius muscle of aged rats of 21-24 months fed with a control or OLE diet equivalent to 25 mg/kg/day for 3 months (n=5 rats/group). (H-J) Mitochondrial respiration after 4 weeks (H), treadmill exercise performance after 2 weeks (I) and muscle weight after 4 weeks of feeding with control or OLE diet equivalent to 50mg/kg/day in 24 month-old mice. (H) Mitochondrial respiratory capacity measured via oxygen consumption rate (OCR) normalized to mean calcein fluorescence in isolated FDB myofibers (n=10 samples / group). (I) Exercise performance assessed in mice fed for 2 weeks with control or OLE diet (n=4 mice/group). (K) Model of Oleuropein-dependent activation of mitochondria via MCU and protection from muscle decline during aging. In all bar graphs, data are presented as mean ± SEM with *p < 0.05 using a two-tailed unpaired T-test.

Taken together, these results demonstrate that oleuropein overcomes MCUR1 decline during aging to rescue MCU activity and prevent bioenergetic and functional performance impairments of aged skeletal muscle across species (**Figs. 6K and S8**).

## DISCUSSION

Targeting mitochondria to boost bioenergetics and energy production has been an area of intense investigation over the past decade given the major role of mitochondrial health in adaptations to lifestyle and environment, and the prominent role of mitochondrial dysfunction in many chronic and genetic diseases. In the present study, we have identified the natural polyphenol Oleuropein as a potent dietary nutrient that directly stimulates energy production in healthy and pathological mitochondria by stimulating mitochondrial respiration and ATP production via mtCa^2+^ uptake. Mechanistically, Oleuropein and its deglycosylated metabolite bind to the MICU1 regulatory subunit of MCU and stimulate the uptake of Ca^2+^ in mitochondria, thereby transiently increasing mitochondrial Ca^2+^ concentrations at physiological levels and activating dephosphorylation of PDH via the Ca^2+^-dependent phosphatase PDP. Other mitochondrial therapeutic strategies have translated to the clinics based on pathological impairments of molecular pathways and metabolic processes, and thereby restoration or stabilization of mitochondrial activity via indirect mechanisms (Singh et al., 2021). Elamipretide (SS-31) protects against mitochondrial dysfunction in genetic mitochondrial diseases by stabilizing the respiratory chain (Chavez et al., 2020; Karaa et al., 2018). Urolithin-A is a natural postbiotic that increases muscle strength in older people by restoring youthful mitophagy (Andreux et al., 2019; Liu et al., 2022; Singh et al., 2022). Finally, NAD^+^ precursors mitigate pathological progression in aging or genetic mitochondrial diseases by replenishing the critical metabolic cofactor NAD^+^ (Lapatto et al., 2023; Martens et al., 2018; Pirinen et al., 2020; Yoshino et al., 2021). In contrast to these adaptive effects, activation of mtCa^2+^ uptake by Oleuropein is direct and occurs minutes after cellular exposure, translating into rapid activation of PDH activity and mitochondrial respiration 2 hours after oral intake. Importantly, Oleuropein can also boost the metabolism of healthy mitochondria via a direct mechanism that acutely stimulates TCA cycle activity. Thus, oleuropein is a unique nutritional solution for mitochondrial therapeutics and the first direct nutritional activator of mitochondrial bioenergetic fluxes.

Oleuropein and its metabolite hydroxytyrosol are enriched in Mediterranean diets with high olive and olive oil consumption (Bulotta et al., 2014; Omar, 2010). Their anti-oxidant and anti-inflammatory properties have been shown to protect from cardiovascular and metabolic diseases and musculoskeletal diseases (Barbaro et al., 2014; Bulotta et al., 2014; Horcajada et al., 2022), and are believed to contribute to the beneficial effects of health-protective Mediterranean diets through molecular mechanisms that remain elusive. Oleuropein is the most abundant polyphenol in olive leaves (Clewell et al., 2016), which are traditionally consumed as infusion or extracts for health benefits in Mediterranean countries and have a history of safe human consumption. Based on this traditional knowledge and recent studies on the health benefits of Oleuropein (Barbaro et al., 2014; Bulotta et al., 2014; Nasrallah et al., 2020), olive leaf extracts with high oleuropein content have been developed for human consumption with documented toxicological studies supporting the safety of oral use in humans (Clewell et al., 2016; Guex et al., 2018). Our results demonstrate that the direct activation of mitochondrial bioenergetics by Oleuropein boosts oxidative metabolism in young skeletal muscle and thereby protects from muscle fatigue and stimulates endurance exercise performance in healthy conditions. Importantly, the action of Oleuropein on mitochondrial activity and muscle performance is both very fast and very specific as KO of the MICU1 subunit that directly binds the molecule abrogates its action on mtCa^2+^, and muscle-specific KO of MCU fully blunts its benefits on mitochondrial respiration and exercise performance.

Our work also establishes impaired uptake of mtCa^2+^ via MCU as a novel hallmark of aging that drives mitochondrial dysfunction, alters energy production and contributes to the progression of sarcopenia. The functional capacity to uptake Ca^2+^ in mitochondria is altered during aging of skeletal muscle both in rodent and human models, and impairs PDH activity, mitochondrial respiration and exercise performance. Both in rodents and human, age-related decline of mtCa^2+^ is caused by the downregulation of the MCUR1 subunit of MCU, while the expression of the structural components of the MCU complex (MCU/MCUb, MICU1/2/3) remain normal. Furthermore, MCUR1 expression is low in muscle biopsies of older people with sarcopenia and mitochondrial dysfunction (Migliavacca et al., 2019), and positively associates with muscle mass, muscle strength and quality of life. While MCUR1 was initially described as a bona fide MCU regulator (Mallilankaraman et al., 2012; Vais et al., 2015), current evidence indicates that it may act as a scaffolding protein for MCU (Tomar et al., 2016), and possibly other inner membrane protein complexes, such as cytochrome c oxidase (Paupe et al., 2015). The mechanism of MCUR1-mediated alteration of mtCa^2+^ uptake during aging is however direct as MCUR1 knock-down in primary human myotubes is sufficient to impair mtCa^2+^ uptake and recapitulate the effect of aging.

Consistent with preclinical studies where genetic modulation of mtCa^2+^ and MCU in skeletal muscle regulates oxidative metabolism, mitochondrial energy production and muscle performance (Gherardi et al., 2019; Mammucari et al., 2015), we demonstrate that the age MCU/mtCa^2+^/PDH/OxPhos signaling in skeletal muscle also controls mitochondrial activity and muscle physiology during aging and is a direct cause of sarcopenia. Importantly, Oleuropein can reverse mitochondrial and functional phenotypes of sarcopenia as the Oleuropein-mediated stimulation of mtCa^2+^ uptake capacity via MICU1 is dominant over the age-related decline of MCUR1 that drives impaired mtCa^2+^ uptake in aged muscle. Restoring PDH activity with Oleuropein or with DCA in aged muscle is sufficient to restore mitochondrial bioenergetic potential and oxidative metabolism to improve muscle fatigue and exercise performance. Chronic oleuropein treatment also increases muscle mass in aged muscle, likely by preventing age-related muscle wasting through secondary adaptation to better performance and activity since muscle hypertrophy was not detected in young mice or after acute treatment. Given its history of safe human use, Oleuropein is therefore a promising candidate for the therapeutic management of sarcopenia and other age-related pathologies with mitochondrial dysfunction and impaired mitochondrial energy production. Its MCU-mediated mechanism of action can be complementary to the anabolic effects of protein and signaling via vitamin D, suggesting that it can be beneficial with well aceepted nutritional solutions for sarcopenia (Cruz-Jentoft et al., 2020; Deutz et al., 2014; Morley et al., 2010).

By combining complementary cellular, animal and clinical studies, our work establishes a causal relationship between mtCa^2+^ uptake via the MCU complex, mitochondrial energy production and muscle performance that responds to physiological, pathological and therapeutic stimulation. The modulation of this metabolic pathway is dynamically regulated via acute nutritional signals that directly modulate metabolic fluxes and mitochondrial activity, and gets impaired chronically during human aging and sarcopenia. Acute consumption of the natural polyphenol Oleuropein translates into optimized muscle energy coupling and ergogenic effects on performance and endurance in healthy conditions, while chronic dietary Oleuropein reverses mitochondrial dysfunction, and impaired muscle mass and muscle performances during sarcopenia. Given the history of safe human use of Oleuropein and polyphenol/olive-rich diets, our discovery opens exciting translation to clinical trials and applications for prevention and medical management of sarcopenia and other age-related diseases.

## Acknowledgments

The authors thank Patricia Coulon for contributing to the measurements of mtCa^2+^ in HSMM from old donors. The authors are grateful to Gyorgy Hajnoczky for sharing control and Micu1^−/−^ MEFs.

## Author contributions

Conceptualization: R.R., J.N.F, C.M., U.D.M; Methodology: G.G., A.W., F.B., B.B., G.E.J., A.H., E.M., M.S., L.N., D.B., S.M., B.Bl., R.R., J.N.F., C.M., U.D.M.; Bioinformatics: E.M.; Modeling: M.S., S.M.; Experimental activities: G.G., A.W., F.B., B.B., A.H.; L.N; Data analysis and interpretation: G.G., A.W., F.B., B.B., G.E.J., A.H., E.M., M.S., L.N., D.B., S.M., B.Bl., R.R., J.N.F, C.M., U.D.M. Writing manuscript: R.R., J.N.F, C.M., U.D.M.; Review and editing manuscript: all authors.

## STAR★ METHODS

### KEY RESOURCES TABLE

| REAGENT or RESOURCE | SOURCE | IDENTIFIER |
| --- | --- | --- |
| **Antibodies** |  |  |
| Rabbit monoclonal anti-MCU | Cell Signaling Technology | Cat#14997 |
| Rabbit polyclonal anti-MCUR | Booster | Cat#A08547-1 |
| Mousse monoclonal anti-TOM20 (F-10) | Santa Cruz | Cat#sc-17764 |
| Rabbit polyclonal anti-MICU1 | Sigma Aldrich | Cat#HPA037479 |
| Rabbit polyclonal anti-PDH | Cell Signaling Technology | Cat#2784<br>RRID:AB_2162928 |
| Rabbit polyclonal anti-PDH (phospho S293) | Abcam | Cat#ab92696,<br>RRID:AB_10711672 |
| Rabbit monoclonal anti-TOM20 (D8T4N) | Cell Signaling Technology | Cat#42406,<br>RRID:AB_2687663 |
| Mouse monoclonal anti-GRP 75 (D-9) | Santa Cruz Biotechnology | Cat#sc-133137,<br>RRID:AB_2120468 |
| Anti-Rabbit Detection Module | ProteinSimple | Cat#DM-001 |
| **Chemicals, peptides, and recombinant proteins** |  |  |
| Histamine | Sigma Aldrich | Cat#H7125<br>CAS #: 51-45-6 |
| Caffeine | Sigma Aldrich | Cat#C0750<br>CAS #: 58-08-2 |
| Epibatidine | Sigma Aldrich | Cat#E1145<br>CAS #: 16637-43-2 |
| Oleuropein | Extrasynthese | Cat#0228S<br>CAS #:32619-42-4 |
| Oleuropein Aglycone | Toronto Research Chemicals | Cat#O532945<br>CAS #:31773-95-2 |
| Hydroxytyrosol | Extrasynthese | Cat#4999S<br>CAS #: 10597-60-1 |
| Oligomycin | Sigma Aldrich | Cat#75351 |
| FCCP | Sigma Aldrich | Cat#C2920 |
| Rotenone | Sigma Aldrich | Cat#45656 |
| AntimycinA | Sigma Aldrich | Cat#A8674 |
| Calcein AM | Sigma Aldrich | Cat#C0875<br>CAS#: 154071-48-4 |
| Human Fibronectin | Corning | Cat#356008 |
| PhosSTOP | Merck | Cat#4906845001 |
| Fura-2, AM, cell permeant | Thermo Fisher | Cat#F1221 |
| Gibco™ MEM, GlutaMAX™ Supplement | Thermo Fisher | Cat#41090 |
| GOOD fetal bovine serum - 500 ml | PanBiotech | Cat#P40-37500 |
| Penicillin-Streptomycin (5,000 U/mL) | Thermo Fisher | Cat#15070063 |
| Gibco™ DMEM, high glucose, GlutaMAX™ Supplement, pyruvate | Thermo Fisher | Cat#31966 |
| Gibco™ DMEM, high glucose, HEPES | Thermo Fisher | Cat#42430 |
| Complete, Mini EDTA-free Protease Inhibitor Cocktail | Merck | Cat#11836170001 |
| RIPA lysis and extraction buffer | Thermo Fisher | Cat#89901 |
| Non-Fat Dry Milk | BioRad | Cat#100-04504 |
| PBS 1X PH 7.4 | Thermo Fisher | Cat# 10010023 |
| Skeletal Muscle Cell Growth Medium | Zen-Bio | Cat#SKM-M medium |
| Dulbecco's Modified Eagle Medium F-12 Nutrient Mixture (DMEM/F-12 (1:1) (1x) +GlutaMAX) | Thermo Fisher | Cat#31331028 |
| FBS | Thermo Fisher | Cat#10270106 |
| Heat-inactivated Horse Serum | Thermo Fisher | Cat#26050088 |
| PBS | Thermo Fisher | Cat#10010023 |
| Tyrode's salts | Sigma Aldrich | Cat#T2145 |
| Dulbecco's Modified Eagle's Medium | Sigma Aldrich | Cat#D5030 |
| TRIzol™ Reagent | Invitrogen, Thermo Fischer | Cat#15596026 |
| SuperScript™ III CellsDirect™ cDNA Synthesis Kit | Invitrogen, Thermo Fischer | Cat# 18080300 |
| Master Mix PowerTrack™ SYBR Green | Thermo Fisher | Cat# A46109 |
| Coelenterazine n | Biotium | 10110<br>CAS #: 55779-48-1 |
| Digitonin | Sigma Aldrich | D5628<br>CAS#: 11024-24-1 |
| N-Benzyl-p-toluenesulfonamide | Sigma Aldrich | BL3H160B8B26<br>CAS#: 1576-37-0 |
| Sodium Chloride | Sigma Aldrich | CAS#: 7647-14-5 |
| Magnesium Chloride | Sigma Aldrich | CAS#: 7791-18-6 |
| HEPES | Sigma Aldrich | CAS#: 7365-45-9 |
| Glucose | Sigma Aldrich | CAS#: 50-99-7 |
| Calcium Chloride | Sigma Aldrich | CAS#: 10043-52-4 |
| Potassium Chloride | Sigma Aldrich | CAS#: 7447-40-7 |
| Thapsigargin | Sigma Aldrich | T9033<br>CAS#: 67526-95-8 |
| EGTA | Sigma Aldrich | E3889<br>CAS#: 67-42-5 |
| Collagenase A | Roche | Cat#COLLA-RO |
| Kaempferol | Sigma Aldrich | CAS#: 5020-18-3 |
| Olive leaf extract standardized with 40% oleuropein (OLE) | Bioactor | CAS#: 8060-29-5 |
| Critical commercial assays |  |  |
| BCA protein assay kit | Pierce Thermo Fisher | Cat#23227 |
| 12-230kD Sally Sue Separation Module | Protein Simple | Cat#SM-S001 |
| Pyruvate dehydrogenase (PDH) Enzyme Activity Microplate Assay Kit (ab109902) | Abcam | Cat#ab109920 |
| NuPAGE™ 4 to 12%, Bis-Tris, 1.0–1.5 mm, Mini Protein Gels | Invitrogen | Cat#NP0336BOX |
| Nitrocellulose membranes | BioRad | Cat#1704158 |
| 12-230 kDa Separation Module | Protein Simple | Cat#SM-S001 |
| Experimental models: Cell lines |  |  |
| Human Skeletal Muscle Cells, Donor 7, Lot# 655307 | Lonza | Cat#LZ-CC-2580 |
| Human Skeletal Muscle Cells, Donor 29, Lot # 639629 | Lonza | Cat#LZ-CC-2580 |
| Human Skeletal Muscle Cells, Donor 215, Lot # SKM070512A | Amsbio | Cat#SKB-F |
| Human Skeletal Muscle Cells, Donor 712, Lot # SKM062712B | Amsbio | Cat#SKB-F |
| Human Skeletal Muscle Cells, Donor 211, Lot # SK120211 | Amsbio | Cat#SKB-F |
| Human Skeletal Muscle Cells, Lot# P01064-69M | Cook Myosite | Cat#SK-1111 |
| MEF cells | G. Hajnoczky (MitoCare Center, Dpt. of Pathology, Anatomy and Cell Biology, Thomas Jefferson University, Philadelphia, Pennsylvania 19107, USA) | PMID: 26956930 |
| MEF MICU1-KO cells | G. Hajnoczky (MitoCare Center, Dpt. of Pathology, Anatomy and Cell Biology, Thomas Jefferson University, Philadelphia, Pennsylvania 19107, USA) | PMID: 26956930 |
| C2C12, Muscle Mouse Myoblast | ATCC | Cat# CRL1772<br>Lot# 509793 |
| HeLa | ATCC | Cat# ATCC-CCL-2 |
| Recombinant DNA |  |  |
| Ad5-U6-shMCUR1 | Sirion Biotech | N/A |
| Ad5-CMV-Mito-Aequorin-Mutant(D119A)-polyA-SV40-Neo | Sirion Biotech | N/A |
| Ad5-U6-shRNAscramble | Sirion, Biotech | N/A |
| Ad5-U6-shMCU#1-U6-shMCU#2 | Sirion Biotech | N/A |
| pcDNA3.1-mtGCaMP6f | available upon request | doi:10.1016/j.celrep.2015.01.056 |
| Software and algorithms |  |  |
| GraphPad Prism Software 7.02 | GraphPad |  |
| Fiji | ImageJ | doi: 10.1038/nmeth.2019 |
| Gen5 Microplate Reader and imager software | Biotek |  |
| XF96 instrument | Seahorse Biosciences |  |
| Other |  |  |
| 35 mm Dish, No 1.5 Coverslip | MatTek | Cat#P35G-1.5-20-C |
| 96 Well White Polystyrene Microplate | Corning | Cat#3903 |
| MicroWell™ 384 well optical bottom plates | Nunc® | Cat#64810 |

### RESOURCE AVAILABILITY

#### Lead contact

Further information and requests for resources and reagents should be directed to and will be fulfilled by the lead contact.

#### Materials availability

This study did not generate new unique reagents.

#### Data and code availability

RNA sequencing data from human muscle biopsies of the Singapore sarcopenia study are available at https://www.ncbi.nlm.nih.gov/geo/query/acc.cgi?acc=GSE111016.

### EXPERIMENTAL MODEL AND SUBJECT DETAILS

#### Cell lines and primary human myotubes

HeLa cells (ATCC, #CCL-2^TM^) were maintained in growth medium consisting of DMEM, GlutaMAX supplement Medium (MEM) (Gibco, Thermo Fisher) containing 10 % heat-inactivated FBS, 100 U/mL penicillin and 100 mg/mL streptomycin (Gibco, Thermo Fisher). Mouse Embryonic Fibroblasts (MEF) (MitoCare Center, department of pathology, USA) and C2C12 cells (ATCC, #CRL-1772) were cultured in growth medium comprising DMEM, high glucose, GlutaMAX™ Supplement, pyruvate (Gibco, Thermo Fisher, #31966) containing 10 % heat-inactivated FBS (Gibco, Thermo Fischer, #26050088), 100 U/ml penicillin and 100 mg/ml streptomycin (Thermo Fisher, #15070063). C2C12 cells were differentiated into myotubes using DMEM, high glucose, GlutaMAX™ supplement, pyruvate (1 mM) (Gibco, Thermo Fisher, #31966) supplemented with 1 % heat-inactivated horse serum (Gibco, Thermo Fischer, #26050088), 100 U/ml penicillin and 100 mg/ml streptomycin (Thermo Fisher, #15070063).

Human Skeletal Muscle cells were purchased from Lonza (#CC-2580: 655307, 639629), Amsbio (#SKB-F: SKM070512A; SK120211; SKM062712B), and Cook Myosite (#SK1111: P01477, P01064, P01590) and kept in Skeletal Muscle Cell Growth Medium (Zen-Bio, SKM-M medium). Differentiation of human myoblasts was initiated by adding DMEM/F-12 (1:1) (1x) + GlutaMAX (Gibco, Thermo Fisher, #31331028) containing 2 % heat-inactivated horse serum (Gibco, Thermo Fischer, #26050088), 100 U/ml penicillin and 100 mg/ml streptomycin (Thermo Fisher, #15070063). All cell lines were maintained at 37 °C and 5 % CO_2_. Changes of incubation conditions are indicated in the respective methods.

Primary human myoblasts from adult donors were obtained from Lonza, Amsbio and Cook Myosite after the supplier received informed consent from the donors and after consent was obtained from the Vaud ethics commission for human research (CER–VD) under protocol 281/14.

#### Animals

Young (3-4 months) and aged (24 months) male C57Bl6J wildtype mice were purchased from Charles River or Janvier Labs. Aged Wistar rats aged 19 months were purchased from Janvier labs. Skeletal muscle specific MCU knockout mice (skMCU^−/−^) were previously generated in the lab of Prof. Mammucari as described in (Gherardi et al., 2019). All animal experiments were approved by the internal ethics committee of Nestlé. All rat studies were approved by the Office Vétérinaire. Cantonal du Canton de Vaud, Switzerland and all mouse studies were approved and performed in accordance with the Italian law D. L.vo n_26/2014. All experiments were carried out in accordance with the European Guidelines for the Care and Use of Laboratory Animals and approved. Animals were raised or acclimatized to the conditions of the animal facility, and housed in a ventilated room under a controlled temperature (21 ± 2°C), with a 12:12h light:dark cycle and free access to chow rodent diet and water ad libitum.

### METHOD DETAILS

#### Human skeletal muscle transcriptomics

RNA sequencing data from Singapore sarcopenia study were extensively described in Migliavacca, E., Tay, S.K.H., Patel, H.P. et al. (Migliavacca et al., 2019) and are available at the Gene Expression Omnibus (https://www.ncbi.nlm.nih.gov/geo/) under accession numbers GSE111016. Here we analyzed the expression of genes controlling mtCa^2+^ uptake and extrusion in human muscle biopsies of the vastus lateralis from older people with and without sarcopenia. Briefly, after removing genes with a mean expression lower than 20 reads, data were normalized by the trimmed mean of M-values method as implemented in the edgeR function calcNormFactors (Robinson et al., 2010) and the voomWithQualityWeights function was applied to model the mean-variance relationship and estimate the sample-specific quality weights (Liu et al., 2015). p Values were corrected for multiple testing using the Benjamini– Hochberg method. Protein interaction network was generated with the MCUR1 protein coding gene using STRING version 11.5 (http://string-db.org/), all data sources, a high confidence score of 0.7 and the maximum number of interactors shown in the first shell set to 50. The interaction network was used as input for functional enrichment analysis using Cytoscape (version 3.9.1) to decipher functionally grouped gene ontology using ClueGO.

#### Mitochondrial Ca^2+^ uptake in primary human skeletal muscle myotubes

Human Skeletal Muscle cells (Lonza, #CC2580) were seeded in a previously prepared 96-well plate, coated with human fibronectin (Corning, #356008) at 20 µg/ml for 1 h at RT and washed twice with PBS 1 X (Thermo Fisher, #10010023) before Skeletal Muscle Cell Growth Medium was added together with 12 000 cells/well. After 24 h, knockdown of MCU or MCUR1 was induced by adenoviral infection with MCU shRNA or MCUR1 shRNA (Sirion Biotech, Germany) at MOI 100. Control cells were subjected to adenoviral infection with scrambled shRNA (Sirion Biotech, Germany) at MOI 100 and all are incubated for 48 h. Cells were then differentiated into myotubes by replacing the medium with Dulbecco’s Modified Eagle Medium F-12 Nutrient Mixture (DMEM/F-12 (1:1) (1x) + GlutaMAX) (Thermo Fisher, #31331028) containing 2 % horse serum (Thermo Fischer, #26050088), 100 U/ml penicillin and 100 mg/ml streptomycin (Thermo Fisher, #15070063). After 3 days, cells were subjected to adenoviral infection with mitochondria-targeted aequorin (Sirion Biotech, Germany) at MOI 200 and incubated for 24 h. For experiments with oleuropein-a, myotubes were washed with aequorin buffer and incubated with native coelenterazine at 5 µM for 2 h at RT in the dark before removing coelenterazine and adding a final concentration of 10 µM Oleuropein-a in 1 % DMSO with aequorin buffer and incubating the cells for 20 min at RT in the dark. Control myotubes were treated with DMSO 1% in aequorin buffer. Myotubes were stimulated with 5 mM caffeine.

#### High throughput screening of mitochondrial Ca^2+^

The high throughput screening includes a primary and an orthogonal screen of mtCa^2+^ and a secondary screen of cytosolic Ca^2+^. Mitochondrial and cytosolic Ca^2+^ concentrations were measured in intact HeLa cells by using the luminescent signal of the mitochondrial- and cytosolic-targeted Ca^2+^ sensors aequorin (Pinton et al., 2007). 9 000 cells/well were seeded in 384-well plates (Corning, #3903) and 15 000 cells/well were seeded in 96 well plates (Corning, #64810), in standard growth medium. After 24 h, HeLa cells were infected with 200 MOI (multiplicity of infection) of the adenoviral vector (Sirion Biotech, Germany) carrying either mitochondria-targeted aequorin (Rizzuto et al., 1992) for primary and orthogonal screening or the cytosolic-targeted aequorin (Brini et al., 1995) for counter screening. After 48 h, the medium was removed and the cells were incubated with 5 µM native coelenterazine (Biotium, #10110-1) diluted in aequorin buffer (145 mM NaCl, 1 mM MgCl_2_, 5 mM KCl, 10 mM HEPES, 10 mM glucose, 1 mM CaCl_2_, pH 7.4) for 2 h at RT in the dark. Then, coelenterazine was aspired and pure compounds were added at 10 µM final concentration in DMSO 1 % for 2 h at RT in the dark. Luminescent signal was measured, in basal condition and after stimulation with 100 µM Histamine (Merck, #7125), diluted in aequorin buffer. To calibrate the Ca^2+^ bioluminescence, HeLa cells were semi-permeabilized with 25 µM digitonin (Merck, #11024-24-1) and 10 mM CaCl_2_ (Merck, #21115) in aequorin buffer. Biolumiscence was detected at the plate readers FLIPR TETRAmax (Molecular Device, USA) and Cytation 3 (Biotek, USA) for 384 well format and 96 well plates format, respectively. The screening library was assembled internally from commercially available natural products that have been reported in food in the Dictionary of Food Compounds (Yannai, 2012). This dictionary contains entries describing natural components of food raw materials and food products and is available at https://dfc.chemnetbase.com/faces/chemical/ChemicalSearch.xhtml.

#### Mitochondrial Ca^2+^ uptake in C2C12 cells

C2C12 cells (ATCC, #CRL-1772) were seeded in a 96-well plate with 10 000 cells/well, in respective growth medium. After 24 h, cells were differentiated into myotubes by replacing the medium with Dulbecco’s Modified Eagle Medium GlutaMAX Supplement High Glucose + Pyruvate (Thermo Fisher, #31966) containing 2 % heat-inactivated horse serum (Thermo Fischer, #26050088), 100 U/ml penicillin and 100 mg/ml streptomycin (Thermo Fisher, #15070063). After 3 days of incubation, cells were subjected to adenoviral infection with mitochondria-targeted aequorin (Sirion Biotech, Germany) at MOI 200 and incubated for 24 h. mtCa^2+^ uptake was determined according to the general protocol using 5 mM caffeine (Sigma Aldrich, #C0750) as stimulant to trigger Ca^2+^ release from the sarcoplasmic reticulum.

#### Mitochondrial Ca^2+^ uptake in semi-permeabilized HeLa cells

To measure mtCa^2+^ uptake in semi-permeabilized HeLa (ATCC, #CCL-2^TM^) and MEF cells (Antony et al., 2016) 35 000 cells/well and 15 000 cells/well respectively were seeded in 96 well plates. After 24 h, cells were infected with the mitochondria-targeted aequorin probe as described in the previous section. After 2 h of incubation with native coelenterazine (Biotium, USA, #10110-1) at 5 µM, cells were washed and first treated with a thapsigargin (Sigma Aldrich, #T9033) solution (0.2 µM) containing intracellular medium (IM: 140 mM KCl, 1 mM KH_2_PO_4_, 1 mM MgCl_2_ and 20 mM Na-HEPES) supplemented with 10 mM Glucose and 0.1 mM EGTA (Sigma Aldrich, #E3889) at pH 7.2 for 15 min. 5 min before starting the measurement, a 60 µM digitonin (Sigma Aldrich, #D5628) solution containing DMSO 1 % for control or 10 µM of the selected compound (in 1 % DMSO) for the treated samples, was added. Basal luminescence was measured before stimulating the cells with a 5X calcium solution (IM supplemented with 8 mM Na-Succinate, 4 mM Na-Pyruvate, 4 mM Mg-ATP, 5 mM Mg-EDTA and 3.12.10^−4^M Ca^2+^_[total]_) corresponding to specific concentrations of Ca^2+^ per well. This concentration of free Ca^2+^ reflects the intracellular Ca^2+^ microdomains and promotes mtCa^2+^ uptake which leads to a rise of the luminescent signal. Finally, a 6X calibration solution (final concentration in the well: 140 mM KCl and 10 mM CaCl_2_) was added to calibrate the signal (Montero et al., 2004). Next, the slope values of mtCa^2+^ uptake were analyzed using linear regression. Peak values during mtCa^2+^ uptake represent the maximal speed of Ca^2+^ uptake.

#### Mitochondrial respiration in primary human myotubes

To measure the effect of oleuropein aglycone on oxygen consumption rate (OCR) in control and MCU-knockdown human myotubes, 8 000 cells/well were seeded in a respective Seahorse XF96 Cell Culture Microplate (Agilent, #101085-004) previously coated with fibronectin (Corning, #356008) at 20 mg/ml for 1 h at RT and washed twice with PBS 1X (Thermo Fisher, # 10010023) before adding the cells and incubating them overnight. The next day, cells were infected with 100 MOI of adenoviral MCU shRNA (Sirion Biotech, Germany) to induce knockdown of MCU. Control cells were infected with 100 MOI of scrambled shRNA (Sirion Biotech, Germany). After 48 h, cells were differentiated into myotubes by exchanging the medium with DMEM/F-12 (1:1) (1x) + GlutaMAX (Thermo Fisher, #31331028) containing 2 % horse serum (Thermo Fisher, #31331028), 100 U/ml penicillin and 100 mg/ml streptomycin (Thermo Fisher, #15070063). After 72 h, cells were washed with KRBH and incubated with 10 µM oleuropein-a (Toronto Chemicals, #O532945) in 1 % DMSO for 20 min at 37 °C without extra CO_2_. For the acquisition, basal respiration was recorded before injecting caffeine (Sigma Aldrich, #C0750) at final concentration 5 mM to stimulate the myotubes followed by 2.5 µg/ml oligomycin (oligo, Sigma Aldrich, #75351) and 2 µM rotenone (rot, Sigma Aldrich, #45656) plus 2 µg/ml antimycin A (AntiA, Sigma Aldrich, #A8674). Basal respiration of control and MCU-kd myotubes was used for normalization and expressed in % to see the effect of oleuropein-a on OCR.

#### Ex vivo muscle force and fatigue

EDL (Extensor digitorum longus) muscles were dissected from tendon to tendon under a stereomicroscope and mounted between a force transducer (KG Scientific Instruments, Heidelberg, Germany) in a small chamber in which oxygenated Krebs solution was continuously circulated and temperature maintained at 25°C. The stimulation conditions were optimized, and the length of the muscle was increased until force development during a 90Hz stimulation was maximal. The force-frequency relationship was recorded by performing muscle activation at increasing frequencies every 15 seconds, starting with a single pulse (twitch), to trains of stimuli at various rates producing unfused or fused tetani (20 Hz, 40Hz, 55Hz, 75Hz, 100Hz, 150Hz). Force was normalized for muscle weight. Oleuropein was added to the medium at a final concentration of 10 µM after the measurement of the first force-frequency relationship. After one hour, we performed a fatigue protocol which consisted of 5 tetanic contractions (100Hz) with a duration of 300ms repeated every second. Fatigue was determined as the force reduction relative to the initial force.

#### In vivo treatments with Oleuropein

Animals were treated orally with an Oleuropein-rich olive leaf extract (OLE) standardized with 40% of Oleuropein as previously reported (Polia et al., 2022). Acute single dose administration was performed by an oral gavage of 50mg/kg OLE dissolved in water, or vehicle, and skeletal muscle was collected after euthanasia 2 to 8h post gavage. Chronic OLE administration was performed via dietary exposure by feeding animals orally with an OLE-diet delivering an OLE intake of 50mg/kg/day in mice and 25mg/kg/day in rats, both corresponding to a human dose of 250mg/day after allometric scaling. Diets were prepared by SAFE, France using an AIN93 purified rodent chow diet supplemented with OLE at 0.44 g/kg for young mice, 0.69 g/kg for aged mice and 0.65 g/kg for aged rats to deliver the intended daily doses based on body weight and food intake of each cohort. Exercise performance on a treadmill was performed non invasively after 2 weeks and force and sample collection were performed after 4 weeks of OLE treatment. After euthanasia, skeletal muscle was quickly dissected and processed freshly or snap frozen in liquid nitrogen and stored at −80°C.

#### In vivo DNA transfection of skeletal muscle

For measurement of mtCa^2+^ uptake in mice, a plasmid encoding the genetic 4mtGCaMP6f Ca^2+^ sensor targeted to mitochondria was electroporated to FDB muscles. Mice were anesthetized and hyaluronidase solution (2mg/ml) (Sigma) was injected under the hindlimb footpads. After 30 min, 20 μg of plasmid DNA in 20 µl of physiological solution was injected with the same procedure of the hyaluronidase. Then, one gold-plated acupuncture needle was placed under the skin at heel, and a second one at the base of the toes, oriented parallel to each other and perpendicular to the longitudinal axis of the foot and connected to the BTX porator (Harvard apparatus). The muscles were electroporated by applying 20 pulses, 20 ms each, 1 s of interval to yield an electric field of 100 V. Single fibers cultures were carried out 7 days later.

#### In vivo muscle force, fatigue and exercise performance

*In vivo* determination of force and contraction kinetics of gastrocnemius was carried out as shown previously (Blaauw et al., 2009). Briefly, mice were anesthetized by a mixture of Rompum and Zoletil and a small incision was made from the knee to the hip, exposing the sciatic nerve. Before branching of the sciatic nerve, Teflon-coated 7 multistranded steel wires (AS 632; Cooner Wire Company, Chatsworth, CA, USA) were implanted with sutures on either side of the sciatic nerve. To avoid recruitment of the ankle dorsal flexors, and therefore significant reduction of torque, the common peroneal nerve was cut. Torque production of the stimulated plantar flexors was measured using a muscle lever system (Model 305C; Aurora Scientific, Aurora, ON, Canada). The force-frequency curves were determined by stepwise increasing stimulation frequency, pausing 30s between stimuli to avoid effects due to fatigue. Fatigue was determined as the force reduction relative to the initial force. The fatigue protocol consisted of six 100 Hz trains of 0.5s once every second. Force was calculated from the torque, assuming an average distance of 2.1 mm between the Achilles tendon insertion and ankle joint. Force was normalized for muscle weight.

Exercise performance was assessed by maximal treadmill running time. For acute concentric exercise studies, mice were acclimated to and trained on a 10° uphill LE8700 treadmill (Harvard apparatus) for 2 days. On day 1 mice ran for 5 min at 8 m min−1 and on day 2 mice ran for 5 min at 8 m min−1 followed by another 5 min at 10 m min−1. On day 3, mice were subjected to a single bout of running starting at the speed of 10 m min−1. 10 min later, the treadmill speed was increased at a rate of 1 m min−1 every 5 min until mice were exhausted. Exhaustion was defined as the point at which mice spent more than 5 s on the electric shocker without attempting to resume running. Total running time was recorded for each mouse.

#### Real time imaging of mitochondrial and cytosolic Ca^2+^ in FDB fibers

Real-time imaging. Muscles were digested in collagenase A (4 mg/ml) (Roche, #COLLA-RO) dissolved in Tyrode’s salt solution (pH 7.4) (Sigma Aldrich, # T2145) containing 10 % fetal bovine serum (Pan Biotech, #P40-37500). Single fibers were isolated, plated on laminin-coated glass coverslips and cultured in DMEM with HEPES (Thermo Fisher, #42430), supplemented with 10 % fetal bovine serum (Pan Biotech, #P40-37500), containing 100 U/ml penicillin and 100 mg/ml streptomycin (Thermo Fisher, #15070063). Fibers were maintained in culture at 37 °C with 5 % CO_2_.

mtCa^2+^ measurements. During the experiments, myofibers were maintained in Krebs-Ringer modified buffer (135 mM NaCl, 5 mM KCl, 1 mM MgCl2, 20 mM HEPES, 1 mM MgSO_4_, 0.4 mM KH_2_PO_4_, 1 mM CaCl_2_, 5.5 mM glucose, pH 7.4) containing 0.02 % pluronic acid for 20 min at 37°C and then washed with Krebs-Ringer modified buffer in presence of 75 μM N-benzyl-P-toluenesulfonamide (Sigma Aldrich, #BL3H160B8B26) to avoid the fiber contraction. 30 mM caffeine (Sigma Aldrich, #C0750) was added when indicated to elicit Ca^2+^ release from intracellular stores. Experiments were performed on a Zeiss Axiovert 200 microscope equipped with a 40×/1.3 N.A. PlanFluor objective. Excitation was performed with a DeltaRAM V high-speed monochromator (Photon Technology International) equipped with a 75 W xenon arc lamp. Images were captured with a high-sensitivity Evolve 512 Delta EMCCD (Photometrics). The system is controlled by MetaMorph 7.5 (Molecular Devices) and was assembled by Crisel Instruments.

To measure mtCa^2+^ uptake in young and aged animals, fibers were dissected and loaded with 2 μM mt-fura-2/AM (Bernabei et al., 2022). Images were collected by alternatively exciting the fluorophore at 340 and 380 nm and fluorescence emission recorded through a 515/30 nm band-pass filter (Semrock). Exposure time was set to 100 ms. Acquisition was performed at binning 1 with 200 of EM gain. Image analysis was performed with Fiji distribution of the ImageJ software(Schindelin et al., 2012). Images were background subtracted. Changes in fluorescence (340/380 nm ratio) was expressed as R/R0, where R is the ratio at time t and R0 is the ratio at the beginning of the experiment.

To measure mtCa^2+^ uptake in oleuropein related experiments, FDB fibers were isolated 7 days after *in vivo* transfection with a plasmid encoding 4mtGCaMP6f (Gherardi et al., 2019). Muscles were digested as described above. 4mtGCaMP6f was alternatively excited every second at 475 and 410 nm respectively and images were acquired through an emission filter (535/30 nm) (Chroma). Exposure time was set to 50 ms. Acquisition was performed at binning 1 with 200 of EM gain. Image analysis was performed with Fiji distribution of the ImageJ software (Schindelin et al., 2012). Images were background corrected frame by frame by subtracting the mean pixel value of a cell-free region of interest. Changes in Ca^2+^ levels (475/410 nm fluorescence ratio) were expressed as R/R0, where R is the ratio at time t and R0 is the ratio at the beginning of the experiment. mtCa^2+^ peak was expressed as (R-R0)/R0 and normalized for the control value.

For cytosolic Ca^2+^ measurements, fibers were dissected and loaded with 2 μM fura-2/AM (Thermo Fisher, #F1221) diluted in Krebs-Ringer modified buffer (described above) containing 0.02 % pluronic acid for 20 min at 37 °C and then washed with Krebs-Ringer modified buffer in presence of 75 μM N-benzyl-P-toluenesulfonamide (Sigma Aldrich, #BL3H160B8B26) to avoid the fiber contraction. 30 mM caffeine (Sigma Aldrich, #C0750) was added when indicated to elicit Ca^2+^ release from intracellular stores. Experiments were performed on a Zeiss Axiovert 200 microscope equipped with a 40×/1.3 N.A. PlanFluor objective. Excitation was performed with a DeltaRAM V high-speed monochromator (Photon Technology International) equipped with a 75 W xenon arc lamp. Images were captured with a high-sensitivity Evolve 512 Delta EMCCD (Photometrics). The system is controlled by MetaMorph 7.5 (Molecular Devices) and was assembled by Crisel Instruments. Images were collected by alternatively exciting the fluorophore at 340 and 380 nm and fluorescence emission recorded through a 515/30 nm band-pass filter (Semrock). Exposure time was set to 100 ms. Acquisition was performed at binning 1 with 200 of EM gain. Image analysis was performed with Fiji distribution of the ImageJ software (Schindelin et al., 2012). Images were background subtracted. Changes in fluorescence (340/380 nm ratio) was expressed as R/R0, where R is the ratio at time t and R0 is the ratio at the beginning of the experiment.

#### Mitochondrial respiration in mouse FDB fibers

For measurements of oxygen consumption rate (OCR) from *in vivo* studies, FDB fibers were isolated following digestion in collagenase A (4 mg/ml) (Roche, #COLLA-RO) dissolved in Tyrode’s salt solution (pH 7.4) (Sigma Aldrich, #T2145) containing 10 % fetal bovine serum (Pan Biotech, # P40-37500). Single fibers were isolated, plated on laminin-coated XF24 microplate wells and cultured in DMEM (Sigma Aldrich, # D5030), supplemented with 1 mM NaPyr, 5 mM glucose, 33 mM NaCl, 15 mg phenol red, 25 mM HEPES, 1 mM of L-Glutamine. Fibers were maintained for 2h in culture at 37 °C in 5 % CO_2_.

To measure exogenous FA utilization, the fibers were cultured in DMEM (Sigma Aldrich, #D5030), supplemented with 0.1 mM NaPyr, 5 mM glucose, 33 mM NaCl, 15 mg phenol red, 25 mM HEPES, 1 mM of L-Glutamine, 0.5 mM carnitine and 100 μM palmitate:BSA. Fibers were maintained for 2 h in culture at 37 °C in 5 % CO_2_.

The rate of oxygen consumption was assessed in real-time with the XFe24 Extracellular Flux Analyzer (Agilent), which allows to measure OCR changes after up to four sequential additions of compounds. Fibers were plated as reported above. A titration with the uncoupler FCCP (Sigma Aldrich, #C2920) was performed, in order to utilize the FCCP concentration (0.6 μM) that maximally increases OCR.

The results were normalized for the fluorescence of calcein (Sigma-Aldrich, #C0875). Fibers were loaded with 2 μM calcein for 30 min. Fluorescence was measured using a Perkin Elmer EnVision plate reader in well scan mode using 480/20 nm filter for excitation and 535/20 nm filter for emission.

#### PDH activity

PDH activity was measured in freshly isolated TA muscles with the Pyruvate Dehydrogenase Enzyme Activity Microplate Assay Kit (Abcam, #ab109920) according to manufacturer’s instructions.

#### RNA extraction, reverse transcription, and qPCR

Total RNA was extracted from mouse tissues through mechanical tissue homogenization in TRIzol™ reagent (Thermo Fisher, #15596026), following manufacturer instructions. The RNA was quantified with Nanodrop (Thermo Fisher) and 1 μg of total RNA of each samples was retro-transcribed using the cDNA synthesis kit SuperScript II (Thermo Fisher, #18080300). Oligo(dT)12-18 primer (Thermo Fisher Scientific) were used as a primer for first stand cDNA synthesis with reverse transcriptase. The obtained cDNA was analyzed by Real-Time PCR using the QuantStudio5 Real-Time PCR System thermocycler and the SYBR green chemistry (Thermo Fisher, # A46109). The primers were designed and analyzed with Primer3 (Rozen and Skaletsky, 2000).The efficiency of all primers was between 90 % and 110 %. The housekeeping genes GAPDH was used as internal controls for cDNA normalization. For quantification, expression levels were calculated by using the 2^−ΔΔCt^ method.

Real-time PCR primer sequences were as follows:

MCU:

Fw 5’-AAAGGAGCCAAAAAGTCACG-3’

Rv 5’-AACGGCGTGAGTTACAAACA-3’

MCUb:

Fw 5’-GGCAGTGAAATTCCAGCTTCA-3’

Rv 5’-CGCTCTCGTCTCTTCTGGATC-3’

EMRE:

Fw 5’-GGACTCTGGGCTCTTGTCAC-3’

Rv 5’-AGAACTTCGCTGCTCTGCTT-3’

MICU1 splice variant specific (NM_144822):

Fw 5’-GCGCTTTGATGGAAAGAAAATTGC-3’

Rv 5’-TGTCTACCTCTCCGTCTCCA-3’

MICU1.1 splice variant specific (NM_001291443):

Fw 5’-CTTTGATGGAAAGGAGTTCTGGC-3’

Rv 5’-CCTCCATGTCTACCTCTCCGT-3’ MICU2:

Fw 5’-TGGAGCACGACGGAGAGTAT-3’

Rv 5’-GCCAGCTTCTTGACCAGTGT-3’

MCUR1:

Fw 5’AGTTTTCAGCCCTCAGAGC-3’

Rv 5’-GGACTTTGGTCACTTCATCCATCA-3’

GAPDH:

Fw 5’-CACCATCTTCCAGGAGCGAG-3’

Rv 5’-CCTTCTCCATGGTGGTGAAGAC-3’

#### Western blot and antibodies

To determine the expression level of MICU1 protein in control and MICU1-KO MEF cells, cells were washed twice using PBS and scraped from a 175 cm^2^ flask to collect the lysate in 50 ml falcons on ice before centrifuging the cells at 1 500 rpm for 4 min at 4 °C. The supernatant was discarded and cells were resuspended in Mito buffer (300 mM sucrose, 10 mM Hepes, 0.5 mM EGTA, Complete EDTA-free protease inhibitor mixture (Merck, #11836170001, Germany), pH 7.4) and incubated on ice for 30 min. Cells were transferred to a 2 ml glass potter for homogenization and transferred to a 2 ml Eppendorf tube and centrifuged at 600 g for 10 min at 4 °C. Supernatant was collected in a 1.5 ml Eppendorf tube and centrifuged at 6 000 g for 10 min at 4°C to pellet the mitochondria, which were dissolved in 50 µl Mito buffer to measure protein concentration via BCA (Pierce Thermo Fisher, #23227). 20 µg of protein was separated by SDS-polyacrylamide gel electrophoresis using 4-12 % acrylamide gels (Invitrogen, #NP0336BOX) and transferred to PVDF membranes (Mini format 0.2 µm PVDF, Bio-Rad) with a semi-dry blotting protein transfer (Trans Blot Turbo, Bio-Rad). TBS-tween-washed membranes (0.5 M Tris, 1.5 M NaCl, 0.01 % Tween, pH 7.4) were blocked with 5 % BSA (Sigma Aldrich, #A2153) for 1 h at RT. The washed membranes were incubated overnight with the primary antibodies MICU1 1:400 (Sigma Life Science, #HPA037479) or TOM20 1:1000 (Cell Signaling, #42406) diluted in 1 % BSA at 4 °C. After 24 h, the washed membranes were incubated for 1 h at RT with IRDye 800 CW goat anti rabbit IgG (LI-COR Biosciences, U.S.) 1:15000 diluted in 1 % BSA. Membranes were scanned on the Odyssey scanner (LI-COR Biosciences, USA).

To determine MCU and MCUR1 protein levels in primary human myotubes, cells were lysed with RIPA buffer (ThermoFischer, #89901) supplemented with Complete EDTA-free protease inhibitor mixture (Merck, #11836170001) and phosSTOP^TM^ (Merck, #4906845001). Protein concentrations were measured via BCA. For Western blotting detection, the Sally Sue automated capillary blotting system (ProteinSimple, San Jose, CA, USA) was applied. Samples were prepared according to the manufacturer’s protocol using the Anti-Rabbit Detection Module (ProteinSimple, #DM-001) and the 12-230 kDa Sally Sue Separation Module (ProteinSimple, #SM-S001). The loading concentration was adjusted to 0.2 µg/µl using RIPA buffer. Primary antibodies used were MCU 1:30 (Cell Signaling, #14997), MCUR1 1:30 (Booster, #A08547-1) and TOM20 1:100 (Cell Signaling, #42406) diluted in the manufactorer’s antibody diluent (ProteinSimple).

To monitor protein levels in mouse skeletal muscles, frozen muscles were pulverized by means of Qiagen Tissue Lyser and protein extracts were prepared in an appropriate buffer containing 50 mM Tris pH 7.5, 150 mM NaCl, 5 mM MgCl_2_, 1 mM DTT, 10 % glycerol, 2 % SDS, 1 % Triton X-100, Complete EDTA-free protease inhibitor mixture (Roche), 1 mM PMSF, 1 mM NaVO3, 5 mM NaF and 3 mM β-glycerophosphate. 40 μg of total proteins were loaded, according to BCA quantification. Proteins were separated by SDS-PAGE electrophoresis, in commercial 4-12 % acrylamide gels (Thermo Fisher, # NP0336BOX) and transferred onto nitrocellulose membranes (Thermo Fisher, #1704158) by semi-dry electrophoretic transfer. Blots were blocked 1 h at RT with 5 % non-fat dry milk (Bio-Rad, #100-04504) in TBS-tween (0.5M Tris, 1.5M NaCl, 0.01 % Tween) solution and incubated at 4 °C with primary antibodies. Secondary antibodies were incubated 1 hr at RT. The following primary antibodies were used: MCU 1:1000 (Merck), phosphoPDH Ser293 1:5000 (Abcam, #ab92696), PDH 1:1000 (Cell Signaling, #2784), GRP75 1:1000 (Santa Cruz, #sc-133137), TOM20 1:1000 (Santa Cruz, #sc-17764).

#### Modeling

3D-model of human MICU1 (UNIPROT entry: Q9BPX6) was obtained from the experimentally solved structure of Ca^2+^-free MICU1 in its hexameric form (PDB ID: 4NSC) (Wang et al., 2014). The strategy adopted to prepare the protein model and the identification of the binding site was previously reported (Di Marco et al., 2020). The ligands were obtained from Pubchem (PubChem CID: 56842347; 5281544) and their partial charge calculated after semi-empirical (PM6) energy minimization using MOE 2018 (Stewart, J.J.P., 2012, MOPAC2012, version 202; http://OpenMOPAC2012.net, Stewart Computational Chemistry). Molecular docking studies were performed with plants1.2 coupled to chemPLP scoring function (Korb et al., 2009) defining as binding site a sphere placed on the model center of the mass and using a radius of 14 Å. For each run 20 output conformations were generated and analyzed by visual inspection.

The ligand-protein complexes were prepared for MD simulations with AmberTools14 (Maier et al., 2015), assigning Gasteiger charges and General Amber Force Field (GAFF) parameters to the ligands and Amber14 partial charges and parameters to the proteins. Each system was solvated with explicit waters (TIP3P model) resulting in a tetragonal box with boundaries at least 11 Å far from any atom of the complex.

Each system was neutralized adding Na^+^/Cl^−^ ions to a final concentration of 0.1 M. Each system was subjected to 300 steps of conjugate-gradient minimization followed by 100 ps NVE and 500 ps NPT equilibration applying harmonic positional constraints (1 kcal mol^−1^ Å^−2^) on protein and ligands atoms. The pressure was maintained to 1 atm by Berendsen barostat and the temperature to 310 K by a Langevin thermostat. Subsequently, three replicas of 10 ns classical MD simulations in the NVT ensemble were conducted to compute MMGBSA energy and assess RMSD along the trajectory.

All MD simulations were carried out with the ACEMD engine (Harvey et al., 2009), with a time-step of 2fs, by handling the nonbonded long-range Coulomb interactions with the particle mesh Ewald summation method (PME) with a cutoff distance of 9 Å and a switching distance of 7.5 Å.

For the analysis of the docking, Chemplp Score/Norm column indicates the value obtained from a scoring function normalized on the number of heavy atoms, measuring the strength of the interaction. Our protocol was extended by using Molecular Dynamics to monitor the stability of the conformation obtained by the docking protocol (RMSD) and the quality of the interaction (MMGBSA score).

#### Data analysis

Data analysis and statistical tests were performed using GraphPad Prism 7.02. All data were expressed as the mean ± SEM. Samples were statistically analyzed using parametric Student’s *t*-test for two groups and ANOVA for multiple groups. Results were considered significant with alpha p-values lower than 0.05; \**p* < 0.05.

For the analysis of aequorin-based Ca^2+^ data, the raw data from the Cytation3 Imaging Reader (BioTek, Switzerland) and FLIPR tetraMAX (Molecular Devices, USA) were calibrated using the following equation (Bonora et al., 2013).

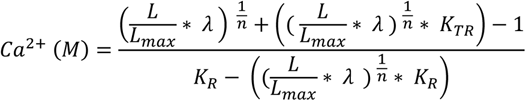

*L* = light intensity at sampling time

*L_max_* = total light emitted at sampling time

*K_R_* = constant for calcium-bound state

*K_TR_* = constant for calcium-unbound state

*λ =* rate constant for aequorin consumption at saturation [Ca^2+^]

*n =* number of Ca^2+^-binding sites

For each calibrated calcium trace, either peak values or slope values using linear regression were analyzed.

