## Supplemental figures and legends for "Mitochondrial Calcium Uptake Declines during Aging and is Directly Activated by Oleuropein to Boost Energy Metabolism and Skeletal Muscle Performance"

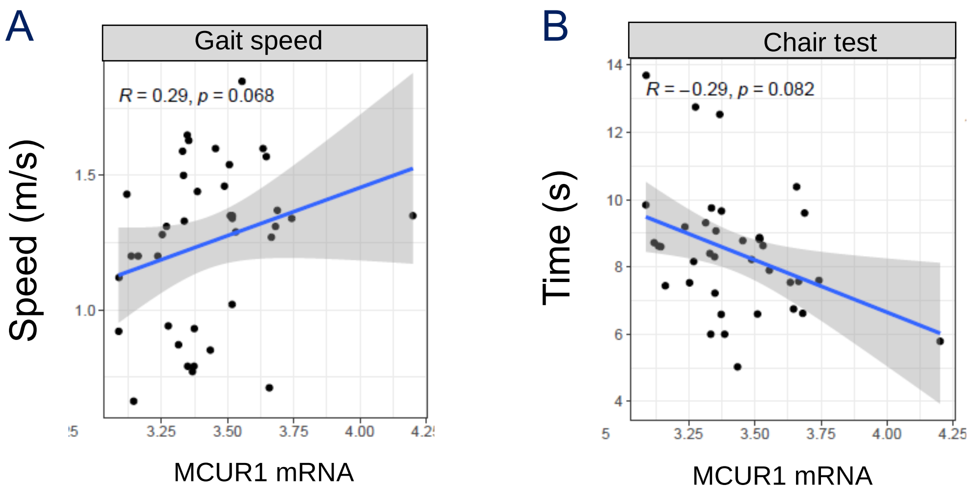


**Figure S1. Association of MCUR-1 to specific muscle features in muscle biopsies of old patients. Related to Figure 1.**
(A, B) Correlation of MCUR1 level with gait speed (A) and chair test (B) in skeletal muscle of old donors.

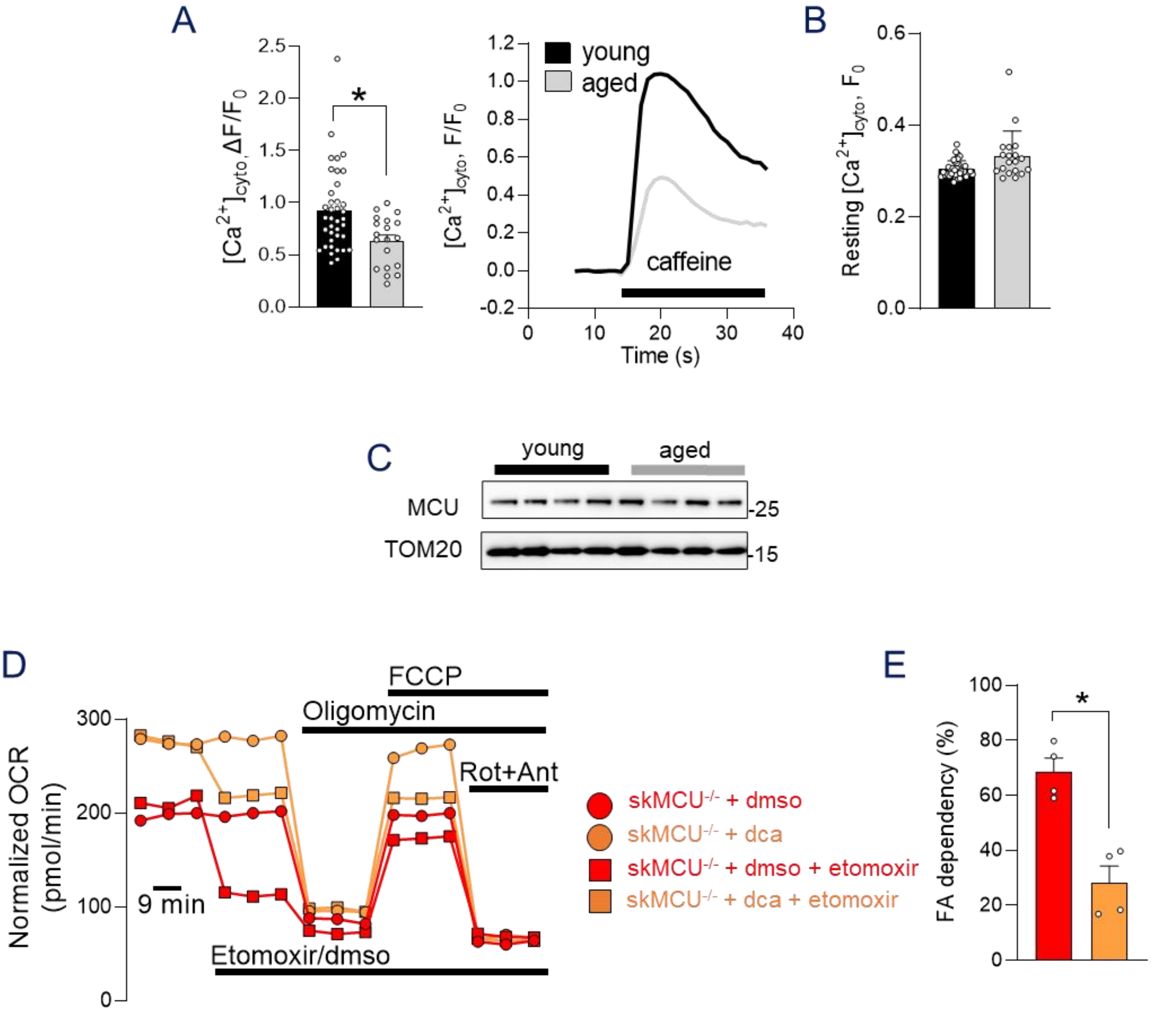


**Figure S2. $Ca^{2+}$ homeostasis is impaired in aged mice. Related to Figure 2.**

(A) Ratiometric measurements of cytosolic $Ca^{2+}$ transients upon caffeine treatment highlighted a reduction in peak cytosolic $[Ca^{2+}]$ of aged FDB myofibers compared to young controls. *$p < 0.05$, t test (two-tailed, unpaired) of >15 fibers per condition. Data are presented as mean ± SEM. Representative traces are reported on the right side.

(B) Resting cytosolic $[Ca^{2+}]$ was increased in aged FDB myofibers compared to young controls. Data are presented as mean ± SEM (>15 fibers per condition).

(C) MCU protein levels do not change in aged muscles compared to young controls. TOM20 was used as loading control.

(D-E) FA-dependent respiration was increased in $skMCU^{-/-}$ FDB myofibers pre-treated with DCA compared to untreated $skMCU^{-/-}$ myofibres. OCR measurements were performed as in Figure 2J. Etomoxir was added to inhibit FA utilization (D). FA dependency was calculated and expressed as percentage of basal OCR (E). Data are normalized on mean calcein fluorescence. *$p < 0.05$, t- test (two-tailed, unpaired) of 4 samples per condition. Data are presented as mean ±SEM.

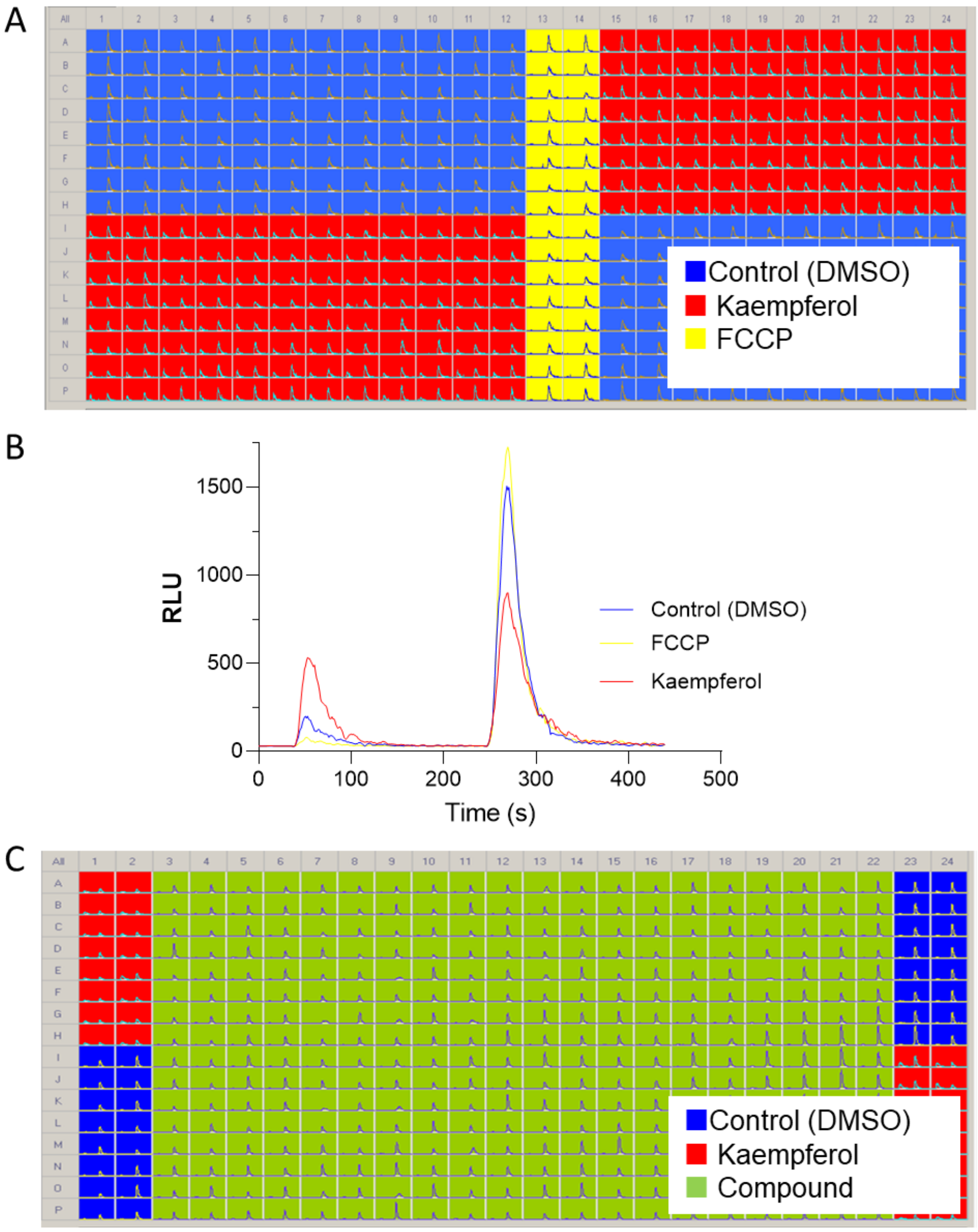


**Figure S3. HTS of mtCa$^{2+}$ and plate design. Related to Fig. 3**

(A) Plate design used for the high-throughput screening of mtCa$^{2+}$. Hela cells were stimulated with 100 mM histamine to evoke mtCa$^{2+}$. Control, DMSO 1% (blue); kaempferol (20 mM , red). FCCP (1 µM) was used as a negative control, to depolarize mitochondria, hampering the driving force of the mtCa$^{2+}$uptake.

(B) mtCa$^{2+}$ traces of Hela cells, stimulated with 100 mM histamine (first peak) followed by addition of digitonin/Ca$^{2+}$ solution to calibrate mtCa$^{2+}$. Control (DMSO 1%, blue), FCCP (1 µM yellow) and kaempferol (20 mM, red).

(C) Plate design used for the high-throughput screening of 5571 compounds. Cells were treated with DMSO (1%, blue) as absolute control, kaempferol (20 mM , red) as positive control and the respective compounds (10 µM, green). Cells were stimulated with 100 mM histamine to evoke mitochondrial Ca$^{2+}$ uptake.

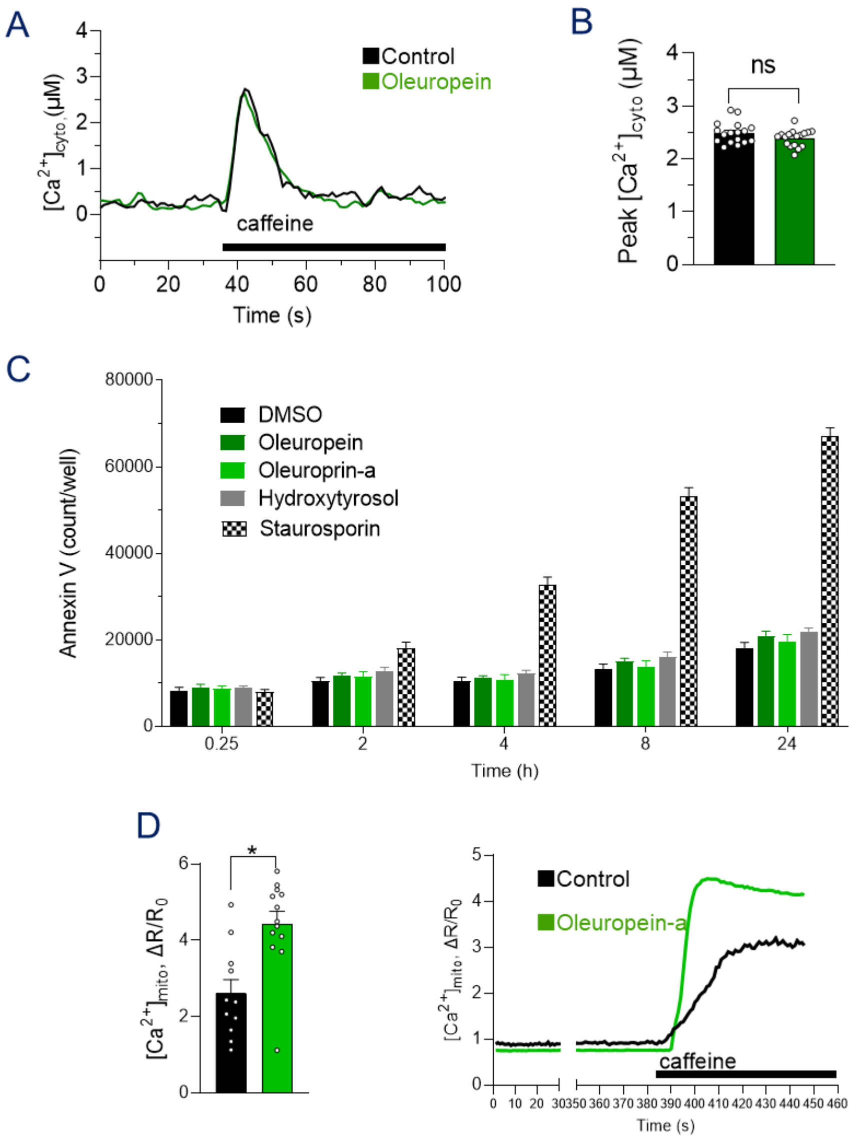


**Figure S4. Oleuropein-a does not affect cytosolic $Ca^{2+}$ levels nor cell death in primary human myotubes. Oleuropein-a increases $mtCa^{2+}$ uptake in muscle fibers. Related to Figure 4.**

(A) Oleuropein-a (10 mM) does not affect cytosolic $Ca^{2+}$ levels after caffeine stimulation (5 mM) in primary human myotubes.

(B) Statistical evaluation of the effect of 10 mM oleuropein-a on cytosolic $Ca^{2+}$ peak, based on data represented in (A). ns: not significant, t-test (two-tailed, unpaired). Data are presented as mean ± SEM of >17 experiments per condition.

(C) Oleuropein and its main metabolites (10 mM) do not affect cell death in primary human myotubes. Control, 1% DMSO. Cell death was measured over a 24 h period as indicated. Staurosporin (100 nM) serves as positive control to increase cell death. Data are presented as mean ±SEM of 6 experiments per condition.

(D) $mtCa^{2+}$ is increased in single isolated mouse FDB fibers treated with oleuropein-a compared to controls. *p < 0.05, t-test (two-tailed, unpaired) of >15 fibers per condition. Data are presented as mean ± SEM. Representative traces are reported on the right side.

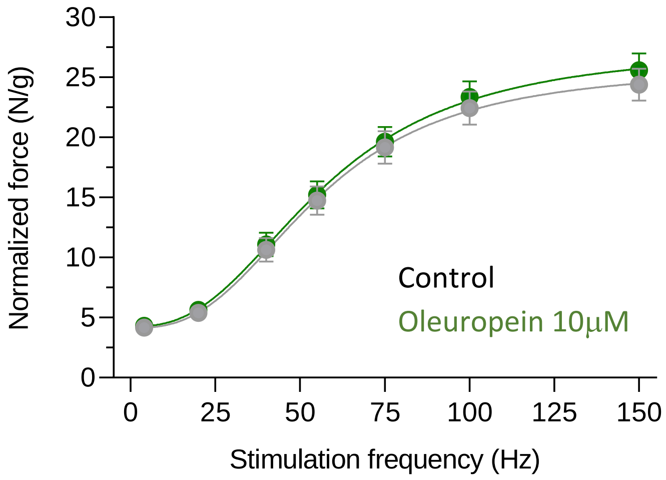


**Figure S5. Oleuropein does not affect normalized force production. Related to Figure 4.** Force-frequency relationship of EDL muscles electrically stimulated in an organ bath *ex-vivo*.The force-frequency relationship was recorded by performing muscle activation at increasing frequencies every 15 seconds. Force was normalized for muscle weight. Oleuropein was added to the medium at a final concentration of 10 µM after the measurement of the first force-frequency relationship. Average of 10 animals per condition.

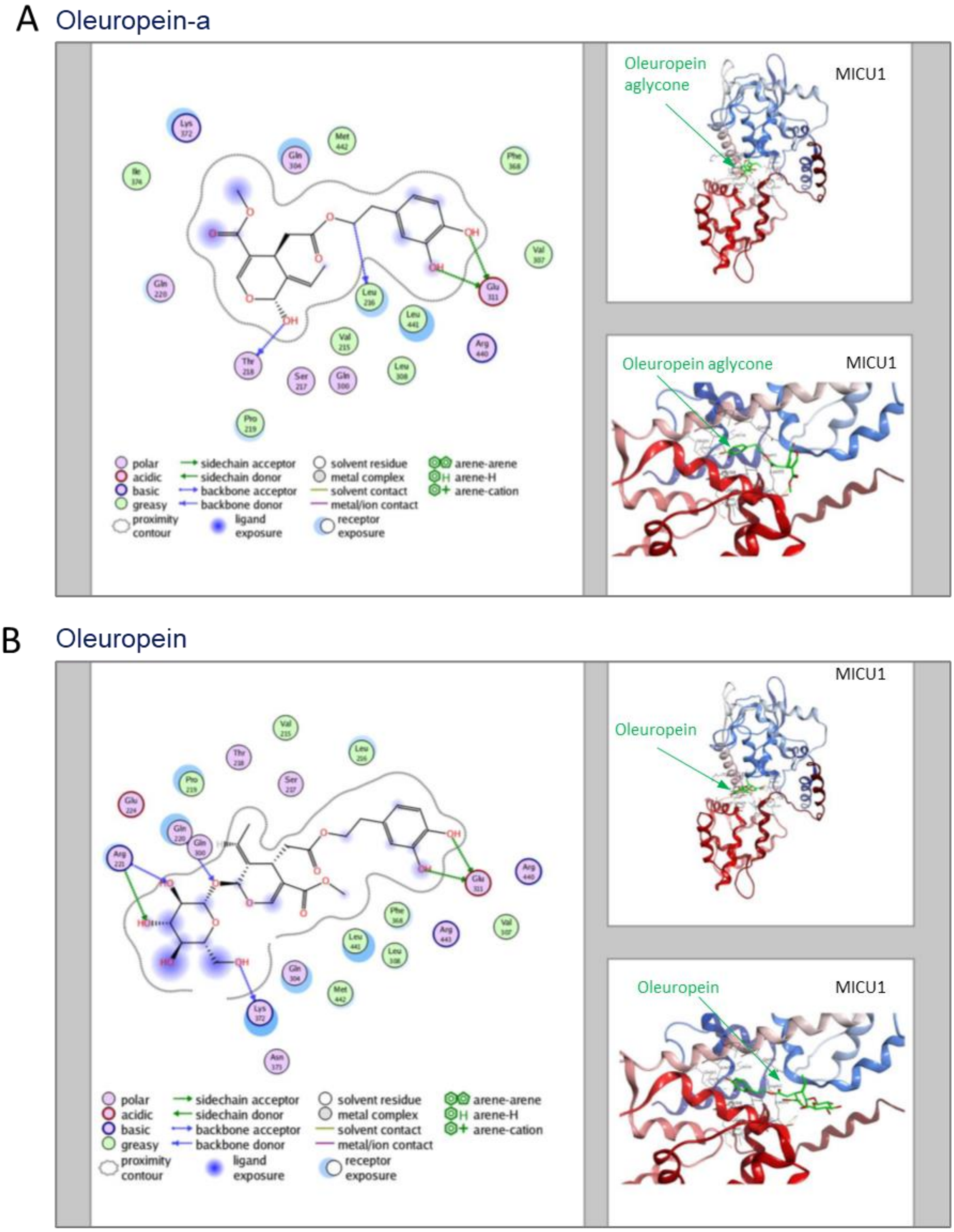


**Figure S6. MICU1-Binding mode of Oleuropein-a and Oleuropein, obtained by docking and followed by MD relaxation. Related to Figure 4.**

The schematic 2D representation of the recognition between MICU1 and the Oleuropein-a in panel A and Oleuropein in panel B. On the right the location of the binding site is reported with two different orientations. The nature of the interactions and the solvent exposure of the interacting residues is color-coded and reported in the legend.

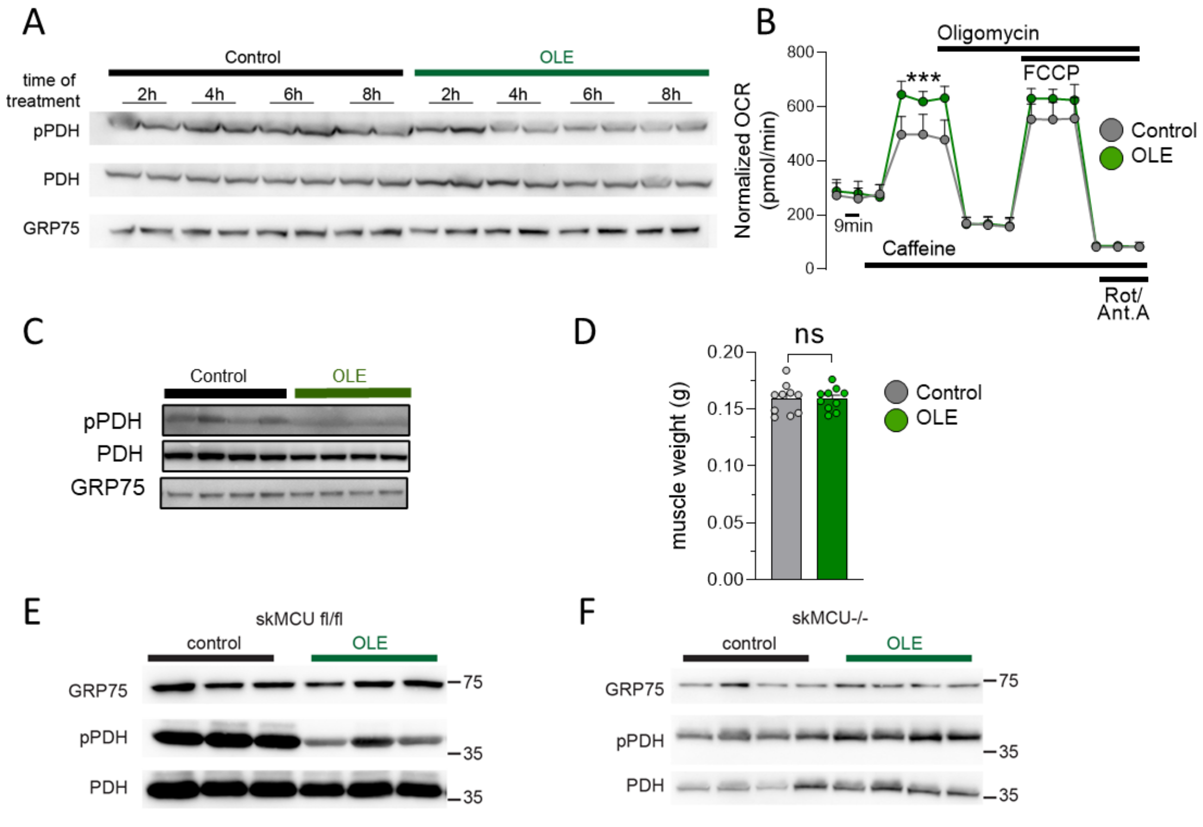


**Figure S7. In vivo chronic OLE treatment in young adult mice. Related to Figure 5.**

(A) Western blotting relative to Figure 5C. GRP75 was used as loading control.

(B) OCR measurements indicate increased respiratory capacity after caffeine stimulation in FDB myofibers of OLE-treated young mice compared to controls. Data are normalized on mean calcein fluorescence. *$p < 0.05$, t- test (two-tailed, unpaired) of 10 samples per condition. Data are presented as mean ±SEM.

(C) Western blotting relative to Figure 5G. GRP75 was used as loading control.

(D) Muscle mass of gastrocnemius muscle of young mice treated with OLE for one month, compared to controls. . *$p < 0.05$, t-test (two-tailed, unpaired). Data are presented as mean ± SEM of 10 muscles per condition.

(E) Phosphorylation levels of PDH were decreased in TA muscles of OLE-treated MCU fl/fl mice compared to controls. GRP75 was used as loading control.

(F) Phosphorylation levels of PDH were unaltered in TA muscles of OLE-treated skMCU$^{-/-}$ mice compared to controls. GRP75 was used as loading control.

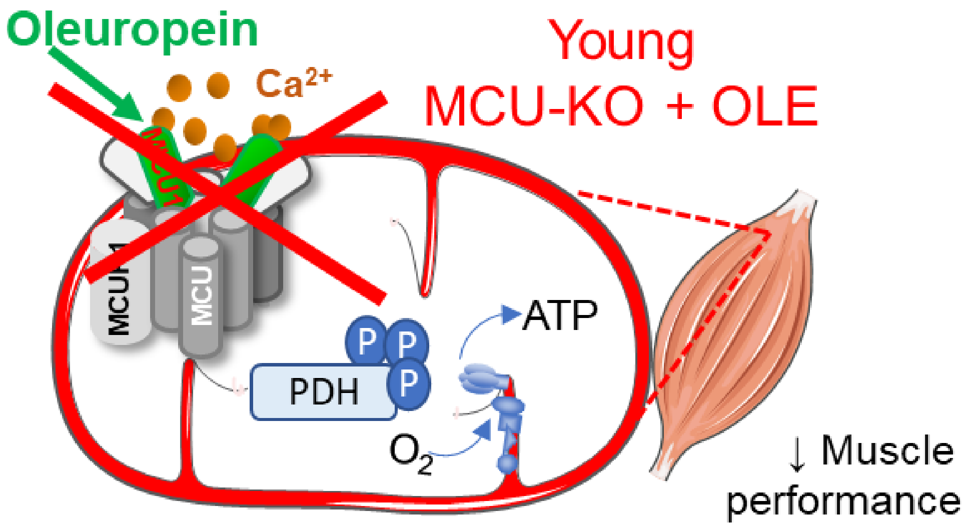


**Figure S8. Model of Oleuropein ineffectiveness in MCU-KO mice. Related to Figure 6.**
